# Cellular porosity in dentin exhibits complex network characteristics with spatio-temporal fluctuations

**DOI:** 10.1101/2025.03.21.644497

**Authors:** Lucas Chatelain, Nicolas Tremblay, Elsa Vennat, Elisabeth Dursun, David Rousseau, Aurélien Gourrier

## Abstract

According to the current hydrodynamic theory, teeth sensitivity is mediated by odontoblast cell processes which can be activated by fluid flow in the pericellular space of bulk dentin. To better understand the possible spatial extent of such phenomena, we investigated the topology and connectivity of dentinal porosity of a healthy human tooth. Using confocal fluorescence microscopy, we modeled the porosity as a spatial graph with edges representing dentinal tubules or lateral branches and nodes defining their connections. A large fraction of porosity channels in crown dentin was found to be interconnected, with 47 % of nodes linked in a single component over a millimetric distance from the dentin-enamel junction (DEJ). However, significant differences in network topology were also observed. A sharp transition in connectivity from 83 % to 43 % occurred at 300 µm from the DEJ, which corresponds to an early stage of tooth formation. This was reflected in all graph metrics investigated, in particular the network resilience which dropped by a factor 2. To test the robustness of our observations, an in-depth analysis of potential remaining biases of the graph extraction was conducted. Most graph metrics considered were found to be within a 10 % precision range from a manually annotated ground truth. However, path metrics, which characterize transport properties, proved very sensitive to network defects. Residual errors were classified in 4 topological classes related to fluorescence staining and confocal detection efficiency, instrumental resolution and image processing. Their relative importance was estimated using statistical and physical graph attack simulations in a broad experimental range. Our modeling thus provides a practical framework to estimate the interpretability of calculated graph metrics for a given experimental microscopy setup and image processing pipeline. Overall, this study shows that dentin porosity exhibits typical characteristics of a complex network and quantitatively emphasize the importance of the smallest lateral branches. Our results could be used to model fluid flow more accurately in order to better understand mechanosensing by odontoblasts in dentin.

## I. Introduction

Odontoblast cells form a dense sensory complex with unmyelinated nerve fibers in the central pulp cavity of the tooth [1,2]. Since odontoblasts are located at the periphery of the pulp cavity, their mechanosensing capacity mostly relies on their ability to probe their spatial environment within the bulk mineralized dentin tissue. To this end, odontoblasts extend a single long cytoplasmic process through dentin inside fluid-filled porosity channels known as tubules which span radially to the dentin-enamel junction (DEJ) [3]. This process has a diameter in the micron range and can be up to 1 mm long inside tubules in human adult molars [4]. It is also known to regularly branch into thinner (0.1-0.5 µm), shorter and more tortuous secondary processes, that either stop at short distances or connect to neighboring cells [3]. This gives rise to smaller secondary porosity channels, termed branches, that, when connecting tubules, allow cell-cell communication [5,6] or activation by pericellular fluid flow at regular intervals along the main process according to the current mechanotransduction theory [7–9]. The overall topology and connectivity of odontoblasts and their processes therefore appear to be key determinants of the mechanosensing capacity of teeth.

Given the apparent importance of lateral branching in the overall dentinal porosity connectivity, Kaye and Herold [5], Holland [10] and Mjör and Nordahl [11] (amongst others) performed detailed histological and statistical characterizations of those secondary processes. However, despite their valuable contributions, a comprehensive understanding of intercellular communication or dentinal fluid flow is still lacking at a large scale. With this in mind, we recently revisited this question focusing on 3D imaging of dentin cellular porosity close to the DEJ in crown dentin [12]. Using confocal fluorescence microscopy, we refined the branching classification proposed by Mjör and Nordahl and revealed that interconnection between odontoblasts via secondary processes may be much more frequent than previously thought, even far from the DEJ. Depending on the spatial extent of secondary processes, odontoblasts could thus form a much more complex network than described so far, which motivates further topological investigations.

Topology analysis based on graph theory forms the basis of complex networks studies across many scientific fields [13]. In biomedical research, various complex networks have been explored, in particular vascular systems in the retina [14,15], liver [16,17], kidney [18,19] and lung [20], as well as pulmonary airways [21,22] and bone porosity networks [23]. It is worthwhile pointing out that all these applications involve transport pathways within organs and could therefore be related to much broader transport network studies [24]. But the field to which graph theory contributed the most is undoubtedly neuroscience. Depending on the imaging or analytical modality used, nodes can represent different brain regions, tissues and even cells [25]. Modeling brain function through the complete set of neurons and their interconnections is currently considered as the ultimate goal of connectomics that aim to decipher structure-function relationships [26,27]. The complexity of such tasks therefore stems from the gigantic number of nodes and edges to be considered, which can lead to extremely challenging mathematics and computational dilemmas.

Inspired by connectomic brain analysis, Weinkamer et al. conducted seminal studies on the osteocyte lacuno-canalicular network (LCN) in bone [23]. Bone tissue closely resembles dentin since it is made of the same structural building block, mineralized collagen fibrils [28–30]. More importantly, osteocytes and odontoblasts both appear to play a key mechanosensory role [31,32]. Using graph representations with nodes as osteocytes and edges as canaliculi, Weinkamer *et al.* laid the ground for advanced numerical 3D fluid flow simulations in the LCN of full osteons sections, thus providing biological insights otherwise inaccessible experimentally [33,34]. They also highlighted small-world properties [35], i.e. any cells in the network that are not directly connected can still be accessed in a small number of steps via nearest neighbors. Despite numerous experimental evidence of tubules interconnections by branches in dentin and its apparent proximity to bone in this respect [36], graph theory hasn’t been used yet to study odontoblast or dentinal porosity connectivity. Such analysis could help elucidate intra- or extra-cellular communication between odontoblasts which could be more complex than currently considered in the literature.

One possible explanation for this gap in our current knowledge of dentin function may be that secondary processes are difficult to image with sufficient resolution over large volumes compatible with connectivity analysis. Transmission electron microscopy (TEM) [37,38] and, to a lesser extent, confocal laser scanning microscopy (CLSM) [4,39,40] provided in-depth insight into cellular components structure, albeit on relatively local scales. To gain a more global perspective of cellular organization, scanning electron microscopy (SEM) was also heavily used to image dentinal porosity, thus providing indirect information on odontoblastic process connectivity [11,41,42]. However, SEM studies were performed on 2D surfaces such that 3D interpretation can only be achieved qualitatively through complex slicing of dentin samples in different orientations. Nevertheless, SEM provided sufficient resolution to capture the finest odontoblastic branches which hasn’t been demonstrated using X-ray tomography so far. As a result, the spatial distribution and connectivity of primary and secondary odontoblast processes are still poorly quantified. Confocal fluorescence microscopy recently provided an indirect but accurate 3D visualization of the odontoblast topology [12]. This technique forms the basis of most connectomic studies performed in the bone field to this date [33–35,43]. Such that all network topology analysis of cells in mineralized tissues published so far rely on the precision of the visualization using confocal fluorescence microscopy and of the graph extraction procedure. Because of the sensitivity of graph analysis to experimental conditions and image processing biases, a rigorous assessment of these factors seems mandatory, though it was rarely performed, if ever.

This study aims to perform a first quantification of odontoblast connectivity in the vicinity of the DEJ, considering dentin porosity as a network. We first describe an image processing pipeline to extract a graph representation of dentinal porosity that can be used for 3D connectivity analysis. In a second step, we examine different classes of graph metrics that best describe the cellular porosity network. We then provide an in-depth estimation of potential residual errors arising from the proposed pipeline and from intrinsic CSLM limitations, thus emphasizing the importance of fine branches that physically bridge tubules during tissue formation. Finally, we analyze the spatial network in relation to well-known histological odontoblast characteristics.

## II. Materials and methods

### II.1. Data acquisition

#### II.1.1. Sample preparation

This study was conducted on a healthy human third molar extracted on 19/02/2018 from a 23 year-old woman with informed oral consent, in accordance with the ethical guidelines laid down by French law (agreement IRB 00006477 and n° DC-2009-927, Cellule Bioéthique DGRI/A5). Immediately after extraction, the tooth was cleaned from oral debris with an ultrasound bath and disinfected in a Chloramine-T (0.5% - 5g/L) solution. It was then fixed in 70 % ethanol for 48 hours and subsequently embedded in an athermal-curing epoxy resin (EPOFIX, Struers) to facilitate handling and cutting. Thin slices of 300 µm in thickness were cut in the mesio-distal plane (Fig.1a) using a water-lubricated diamond saw (Struers E0D15 mounted on a Presi Mecatome T210). The sections were cleaned with water from cutting residues and polished down (Presi Minitech 233) with SiC paper up to P4000 grade (EU grade, Struers) to 200 µm in thickness measured with a digital gauge. Finally, the sample was stained in an 0.02 wt% Rhodamine B glycerol solution for 72 h at room temperature and sealed between two glass cover slides in the same solution.

**Figure 1:**
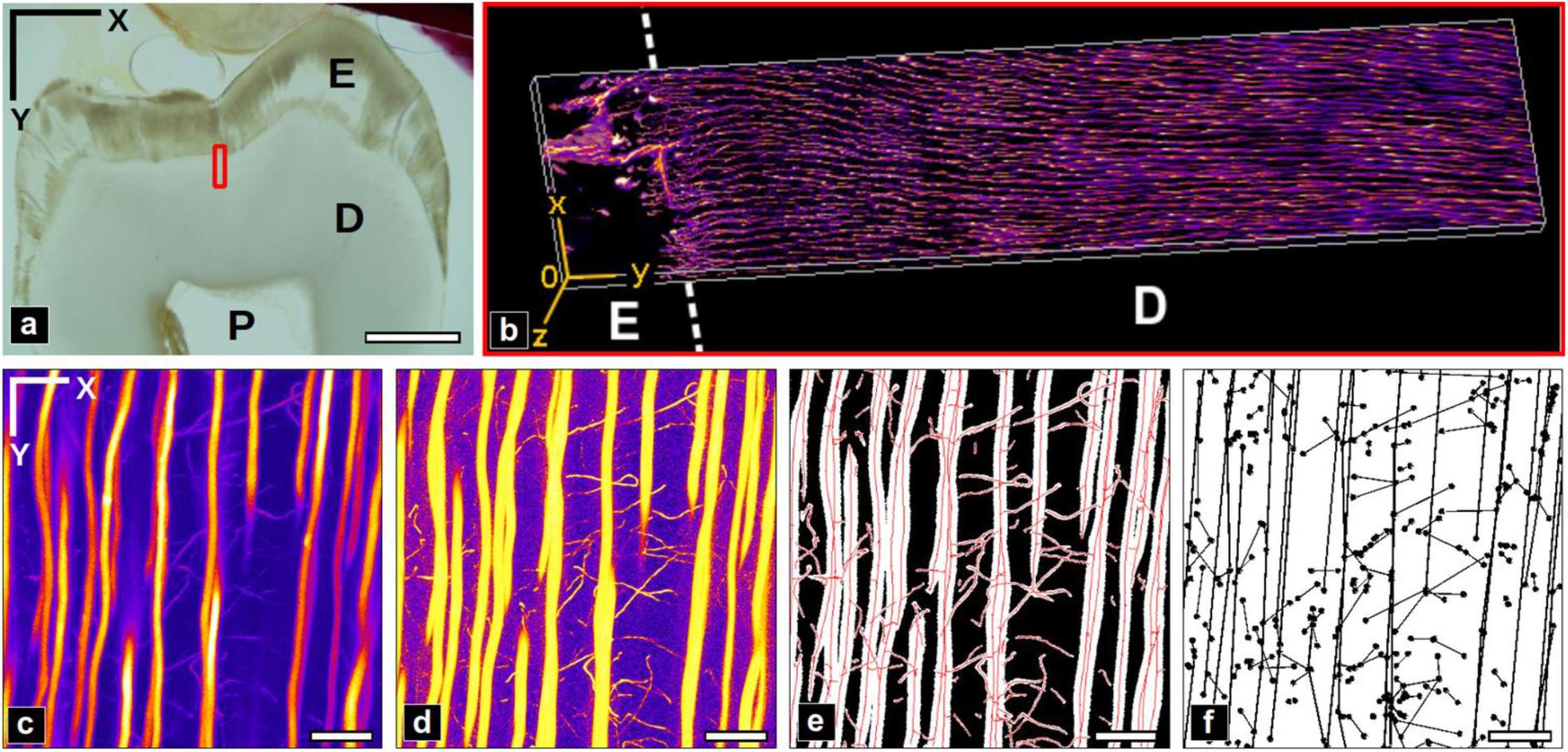
Methods summary. a) Transmission light microscopy of the sample section imaged with confocal laser scanning microscopy (CLSM) in the area indicated by a red rectangle (width: 200 µm, height: 900 µm). E: Enamel, D: Dentin, P: Pulp cavity. Scale bar: 2 mm. b) 3D rendering of the resulting image stack. c-f) Image processing pipeline of the image stack: c) original image, d) vesselness filtering, e) porosity segmentation with its skeleton (red), f) cleaned graph. Displayed images are a maximum intensity projection (MIP) of 51.2×51.2×20 µm^3^ volumes. Scale bar: 10 µm.

#### II.1.2. Confocal fluorescence imaging

Confocal image stacks were acquired in the period 26/02/2018 to 09/03/2018 with a Leica TCS SP8 microscope, using a 40x/1.3NA oil immersion lens. A wavelength of 561 nm (LED laser DPSS 561) was used to excite Rhodamine B fluorescence which was collected in a range of 567-636 nm with a hybrid detector. Acquisitions were performed on the polished surface of crown dentin up to 900 µm from the DEJ towards the pulp (cf Fig.1a-b). The scanning was performed using 26-52 µW laser power with a linear increase in depth to compensate for scattering and absorption with a pixel dwell time of 300 ns and a line average of 3 acquisitions. The imaged volume consisted of a mosaic of 5 overlapping stacks, each consisting of 205.5×205.5 µm^2^ images with a pixel size of 100 nm, collected over 20.65 µm in depth with 350 nm sampling, forming a 934.4×205.5×20.65 µm^3^ volume. The resulting dataset was thus 2055×9344×59 voxels in shape with 100×100×350 nm^3^ voxel size.

### II.2. Processing pipeline: from 3D image stacks to graphs

Three different graphs are considered in this study and described in more details in the following sections:

- the generated graph: a coarse graph automatically generated from the raw image stack following the image processing pipeline.
- the cleaned graph: a semi-automatically generated version in which generic errors from the generated graph after systematically corrected.
- the ground truth (GT) graph: obtained by manual annotation of the data to correct remaining errors.

#### II.2.1. Porosity segmentation

All calculations were performed on a computer equipped with an Intel Xeon 1270 CPU (3.40 GHz 16) and 64 Gb of RAM. Data processing was done using custom Python code (based on the numpy [44], scipy [45] or scikit-image [46] libraries unless specified otherwise). The raw mosaic volume was first cropped to remove the enamel region and the intensity level of the stack of images was adjusted by histogram equalization, to account for the intensity attenuation in depth due to scattering and absorption. The dataset was then resliced in Z (in-depth direction) by a factor of 3.5 (with bicubic interpolation) to obtain isotropic voxels of 100×100×100 nm^3^. The final dataset size was thus 8196×2055×206 voxels.

As dentinal porosity appears in the form of tubes with varying diameter, a morphological multiscale 3D vesselness filter was applied to enhance vessel structures in the imaged volume [47]. The filter is based on the eigenvalues of the Hessian of the image that can help in assessing the structure shapes locally. Tubular features are identified as locations with two strong eigenvalues (in cross-section) and a small one (along the vessel axis). The implementation from Jerman [48] was used with parameters σ (set of scales for the hessian) in the range 1-5 pixels in steps of 0.05 (corresponding to 2.36-11.78 pixels FWHM, or 0.236-1.178 µm) and τ = 0.5. The filtered volumetric porosity was then segmented using hysteresis thresholding with thresholds determined using a 3-class multi-Otsu method. Remaining noise was removed from the final binary porosity mask through a sequence of morphological opening (3 voxel diameter ball footprint), removal of isolated voxels clusters (either black or white, of size less than 256 voxels with 26-connectivity) and final median filtering (3 voxels side length cubic window).

#### II.2.2. Topological graph extraction

The skeleton of the smoothed 3D binary mask was obtained with a topology preserving thinning process (Lee’s method [49]) to reduce the vessels to their centerlines. A multigraph, as defined in the NetworkX library [50] was extracted from the skeleton via the sknw Python library [51]. The generated graphs have nodes at branching or end points and edges representing the vessels. Multiple node and edge attributes were also created in the process. Each node and edge first have an attribute corresponding to the list of associated voxels positions to retain the skeleton information in the graph. Then, the node center of gravity (defined by 1 to 3 skeleton voxels) was added as an attribute. For the edges, a “weight” attribute corresponding to the arc length (sum of the pairwise distances in “pts”) was also created.

#### II.2.3. Graph cleaning

In a first step, the generated graph was filtered to remove undesired artifacts and topological errors. The algorithm developed for this cleaning is detailed in S1 Fig. The terms “artifacts” refer to elements in the graph that don’t correspond to our biological knowledge of dentin porosity. This includes multi-edges (different edges associated to the same pair of nodes), self-loops (edges linked to the same node) and nodes of degree equal to 2 (nodes with two edges). The second step consisted in removing “errors” defined as sets of nodes and edges that do not correspond to real porosity vessels and branching points. The removals of both spurious edges and edge bridges are considered, two types of errors described in more detail below (III.1.). The cleaning of spurious edges concerns the removal of terminal edges (edges with at least one node of degree 1) based on a threshold over the “bulge size” parameter defined in [52]. The cleaning of a specific kind of edge, defined as “edge bridges” in III.3.1, is based on local orientation of the edge and its neighbors (see S1 Fig. for more details). The graph was thus cleaned from “artifacts” and “errors” through several iterations until no changes were detected.

#### II.2.4. Spatial maps of graph metrics

Coarse-grained maps of the graph metrics were obtained by applying a sliding window in the XY plane to define patches of size 25×25×20.06 µm^3^ with a 50% overlap. Within each patch, subgraphs were defined, which contain all nodes and edges of the global graph within the patch. Graph metrics were computed on each subgraph and the resulting values were then stored in a matrix array defining a spatial map, where each pixel correspond to the subgraph position in the sample.

#### II.2.5. Ground truth creation

The graph obtained after the automatic cleaning step was further corrected via a manual annotation in two sub-volumes of interest (ROI) of 1000×1000×206 voxels region. The manually corrected graphs are considered in the following as “ground truth graphs”. The annotation was done with FIJI [53] in the form of markers. 3D image representations where the graph was superposed to the original image stack were used to visually compare the resulting graph with the acquired 3D volume. Annotation markers were then exported and used in Python scripts to apply the modifications. Three types of graph corrections were considered: deletion (node or edge), creation of a missing straight edge, creation of a node and creation of an edge with a broken line path.

#### II.2.6. Graph representation

The software Gephi [54] was used to display the graphs. The Fig.4.b,e representation was obtained using the node positions, and the one for Fig.4.c,f is based on the use of multiple predefined layouts (OpenOrd, Force Atlas and Noverlap).

### II.3. Network analysis

#### II.3.1. Graph metrics

Because graphs representing cellular and porosity networks in bone and teeth are essentially spatial graphs, we use the corresponding terminology for graph metrics proposed by Barthélemy et al. [55], a reference review on the topic. A limited subset of the metrics described by the authors was selected to account for different categories of information.

##### Graph descriptors

***N***: Total number of nodes

***E***: Total number of edges

***⟨ k⟩*** : Mean degree: average number of node neighbors

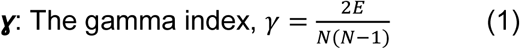

The number of nodes ***N*** and of edges ***E*** are the most fundamental network properties. The mean node degree **<*k*>** and gamma index **ɣ** (Eq. 1) respectively represent the average number of neighbors a node possesses and the probability of having an edge between two randomly selected nodes. Altogether, these metrics provide a basic understanding of the graph’s size and sparsity.

##### Connectivity metrics

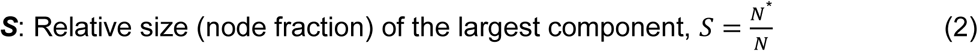

***K***: Number of connected components

***FS***: Fault sensitivity, fraction of edges in the largest component whose removal would break the component in two

where the symbol * denotes a calculation on the largest connected component (in number of nodes) of the graph.

Tubules and branches cut at the borders of the imaged volume are represented by edges which may appear disconnected, giving rise to small connected components. The relative size of the largest component ***S*** (Eq. 2) therefore allows assessing the proportion of nodes forming the connected core of the graph. The number of connected components ***K*** (Eq. 6) is used to evaluate how fragmented the graph is. The robustness of the network can be further explored with the fault sensitivity ***FS*** (Eq. 3), the probability to create a new component when removing an edge at random. Note that this metric was defined as fault tolerance in Barthélemy et al. [55] but we find this terminology misleading since it somewhat represents the “intolerance” to defects.

#### Path-based and distance-based measures

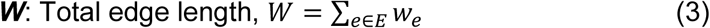

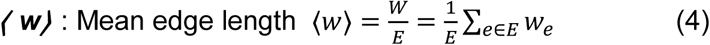

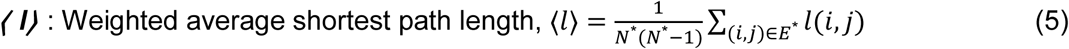

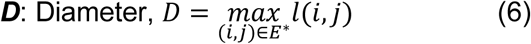

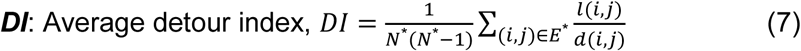

where ***W***, **<*w*>**, **<*l*>** and ***D*** are in µm, with *w_e_* the length of edge e, 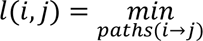|*path*| and *d*(*i*, *j*) respectively the shortest route length and the Euclidean distance between two nodes.

For a spatial graph where edges represent physical paths, the evaluation of weighted distances provide valuable insight to the transport properties of the network. The total edge length ***W*** (Eq. 3) is the global path quantity, while the mean edge length **<*w*>** (Eq. 4) give a basic estimate of the spacing between nodes. The weighted shortest path length ***l(i,j)*** corresponds to the minimum route distance (in µm) between two nodes. The weighted average path length **<*l*>** (Eq. 5) and diameter ***D*** (Eq. 6) (the longest shortest path) yield intuitive measures of node proximity in the graph. In this study, only nodes from the largest component (hence the star symbols) were considered to avoid biases from other smaller components. The average detour index ***DI*** [Eq. 7], which is the mean ratio of the shortest path length (or route length) to the Euclidean distance for each pair of nodes, also provides a valuable measure of the network travel efficiency.

#### II.3.2. Assessing the robustness of the chosen graph metrics

For each identified type of graph error (see section III.3.1 and table 4) and each graph metric, two simulations were performed starting from the ground truth graphs: random modifications, akin to widely performed “graph attacks” [56] and targeted “physics- or biology-informed” modifications which makes use of the spatial graph attributes as described below.

##### Node disconnections

**Physical model**: nodes of degree superior or equal to 3 are selected and their edge of smallest diameter is disconnected by removing a few voxels next to the node (according to the diameter at the node position).
**Random model**: for each node of degree superior or equal to 3, an edge is randomly removed

##### Edge bridges

**Physical model**: edges are added where vessels are close to each other, based on a distance threshold. For every edge voxel, compute the Euclidean distances to all other voxels (from different edges) and keep the shortest. The distances are weighted such that X and Y length are multiplied by 3.5 to be representative of the dimensions of the instrument’s PSF, meaning that bridges tend to be aligned along the Z axis. Bridges are detected as the pairs of voxels with shortest distance, sorted by increasing length and added in this order up to a certain threshold during the simulation.
**Random model**: edges are added between randomly picked edge voxels (previously created bridges excluded).

#### Missing edges

**Physical model**: based on the intensity value of each voxel, edges of the graph are erased where their diameter is below a given threshold, leading to disconnections, splitting in several pieces or complete removal. During the simulation, this threshold is linearly increased from 0 to the maximum diameter value in the graph by steps of 50 nm (half a pixel length).
**Random model**: edges are randomly removed until none remain in the graph.

## III. Results

### III.1. Morphology of the dentinal porosity network

In contrast to the enamel region (cropped for the analysis), where the only visible features are cracks stained by RhB, a dense packing of elongated tubular structures is visible in the confocal fluorescence stack of stained dentin with variable intensity but high contrast (Fig.3). Those correspond to open cellular porosity infiltrated by the fluorescent solution, which was abundantly described in 2D by SEM at higher resolution [57,58] and confocal microscopy [37,39]. The largest bright vessels roughly aligned along the radial DEJ/pulp direction (vertically in Fig.3)correspond to tubules. Thinner and more tortuous branches can also be visualized with high contrast which allows discriminating those that connect tubules from those that simply extend in dentin (Fig.3.f-g). Some branches also extend radially over large distances (Fig.2.h). Rotating the 3D volume by 90° about the X axis (Fig.3.i-j) allows observing the tubules in cross-section, as often done in SEM studies to evaluate tubule density and diameters. Tubules and branches appear elongated along the Z axis because of the axial anisotropic point-spread-function (PSF) of the confocal microscope.

**Figure 2:**
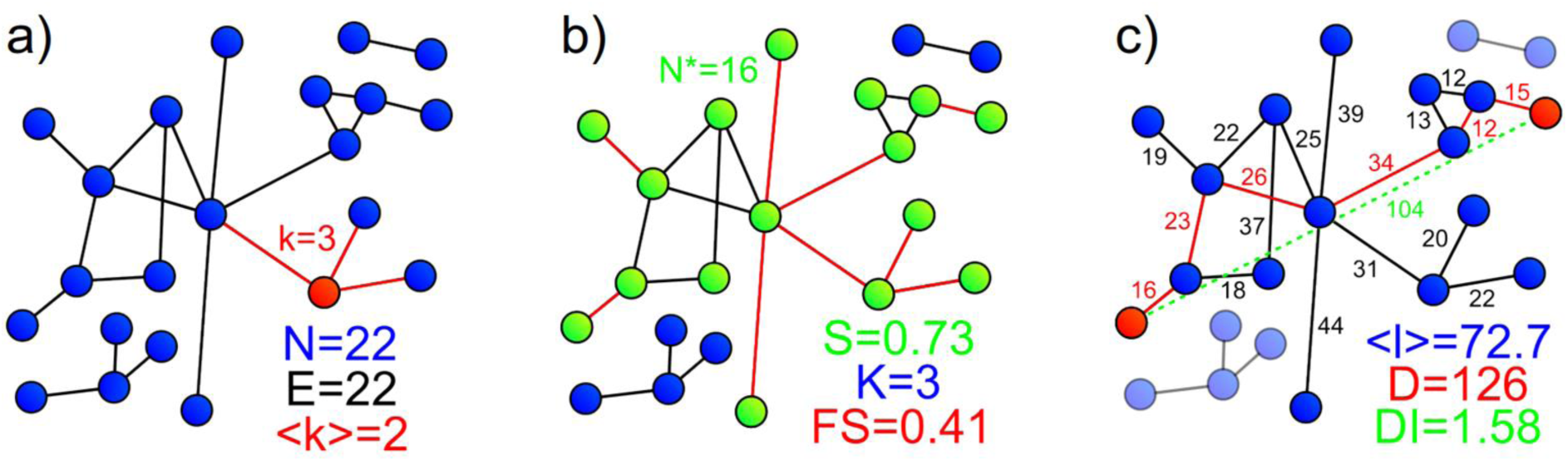
Graph metrics summary. a) **Graph descriptors**. ***k*** is the degree of the node highlighted in red. b) **Connectivity metrics**. Highlighted nodes in green show the largest connected component containing ***N\**** nodes. ***FS*** is the ratio of edges (in red) which removal would lead to disconnections. c) **Distance metrics**. All components but the principal one are faded out to indicate that, in this study, the distance metrics are computed only in the principal component. Displayed numbers correspond to edge lengths. The edges highlighted in red represent the shortest path between two highlighted nodes. The shortest path in this example is the longest in the graph, meaning that its length corresponds to the graph diameter ***D***. The detour index of this specific path is 1.21 (ratio between the route length in red and the Euclidean distance in dashed green). The graph detour index (***DI***) is the average of the detour index of all shortest paths in the principal component.

**Figure 3:**
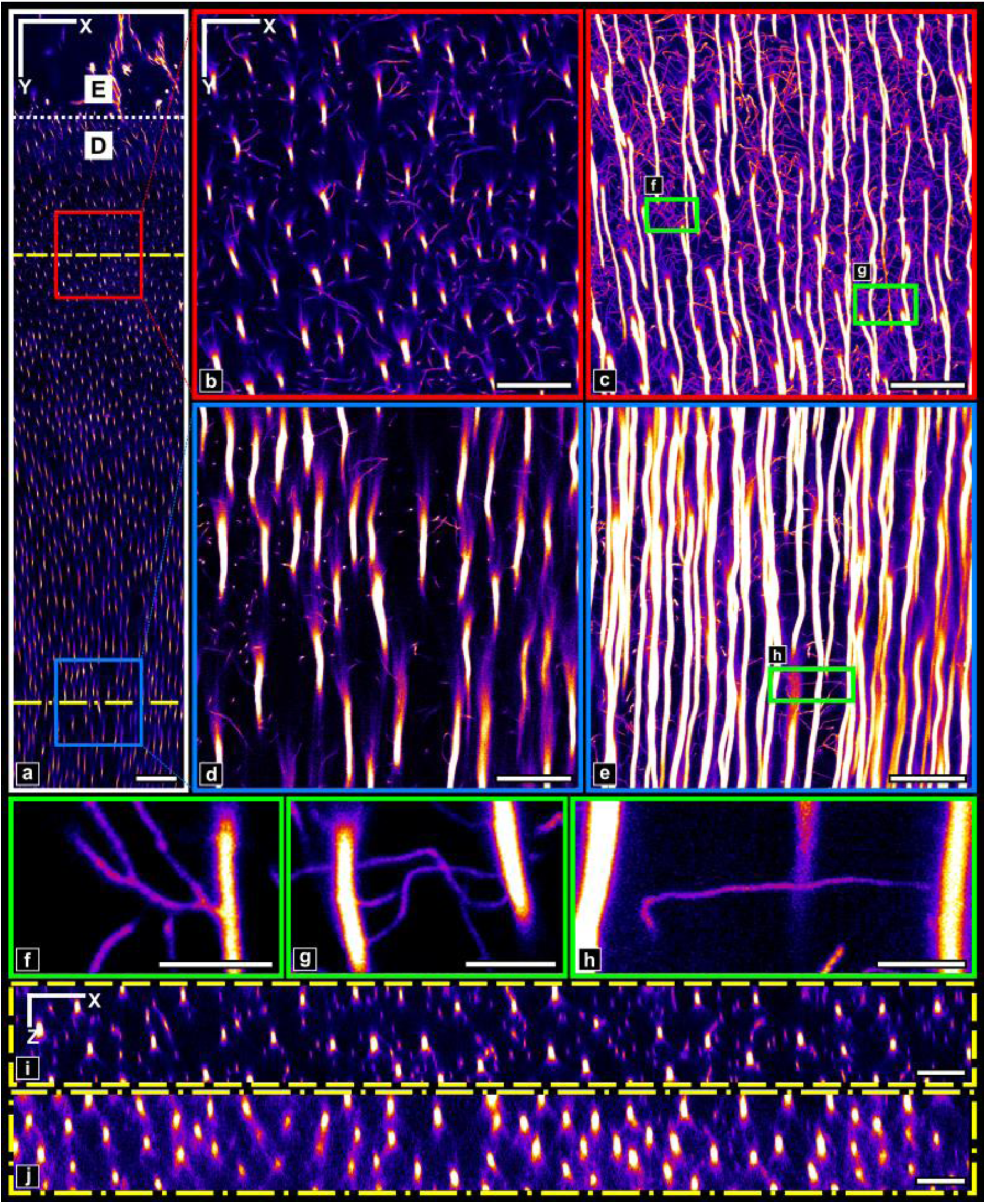
Acquired image stack visualization. a) First slice of the image stack. The red and blue boxes show respectively ROIs 1 and 2 used for further analysis. The dotted white line highlights the different regions, marked by the letters D: dentin and E: Enamel. Yellow dashed lines indicate the location of the orthogonal cuts (in the XZ plane) displayed in f) and g). Scale bar: 50µm. b) ROI 1 first slice. c) ROI 1 maximum intensity projection (MIP). d) ROI 2 first slice. e) ROI 2 MIP. b-e) Scale bar: 10µm. f-h) MIP over a limited number of slices (respectively 10, 9 and 17 slices) in sub-regions of ROIs. Scale bar: 5µm. i) XZ orthogonal cut close to the enamel. j) XZ orthogonal cut far from the enamel. i-j) Scale bar: 10µm. All images were contrasted (different values were used for each ROI) to enhance the visual perception of the structures.

The DEJ chronologically marks the onset of dentinogenesis by odontoblasts. Interestingly, two regions taken near (150 µm, Fig.3.b,c) and far (700 µm, Fig.3.d,e) from the DEJ exhibit distinct topological characteristics. First, tubules appear wavy close to the DEJ and straighter at longer distances. Secondly, the density of tubules increases further from the DEJ, although image sharpness decreases due to optical aberrations. Third, the branches density and contrast seems to decline far from the DEJ. Those observations are in perfect agreement with existing literature (see e.g. [5,11,12,59]).

### III.2. Topological graph characteristics

#### III.2.1 Initial global estimation from extracted graphs

It is well known that many artifacts can result from the graph extraction process [52,60] which typically requires removing nodes and/or edges. However, in the absence of universal graph cleaning rules, corrections ultimately depend on expert user input and their implementation in computing libraries. Furthermore, the impact of nodes and/or edges removal varies for each type of graph metrics. Table 1 summarizes the differences between the generated and corrected cleaned graph of the whole sample volume, based on a set of custom rules described in II.2 and S1 Fig.

**Table 1:** Global graph analysis. The values for different metrics are compared between the generated and cleaned graphs.

| Metric | Full graph |  |
| --- | --- | --- |
|  | Generated | Cleaned |
| $N$ | 92226 | 38755 |
| $E$ | 93621 | 35995 |
| $\langle k \rangle$ | 2.03 | 1.86 |
| $\gamma (\cdot 10^4)$ | 0.22 | 0.48 |
| $\langle w \rangle$ | 2.59 | 6.07 |
| $W (\cdot 10^{-3})$ | 239.5 | 218.6 |
| $\langle l \rangle$ | 265 | 311 |
| $D$ | 1108 | 1017 |
| $DI$ | 2.37 | 2.39 |
| $S$ | 0.56 | 0.50 |
| $K$ | 5727 | 6197 |
| $FS$ | 0.68 | 0.61 |

First, it can be noticed that the cleaned graph is very sparse, as evidenced by the similar number of nodes ***N*** and edges ***E*** (∼ 35 k). This derives from the fact that ∼ 80 % of the nodes have a degree (number of neighbors) of 1 (end of branch) or 3 (tubule/branch connection) (S2 Fig.), which results in low mean degree **<*k*>** ∼ 2 and gamma index ***γ*** values ∼ 5.10^-5^. This implies that, in most cases, only a single branch stems from the main tubule or joins another one at a given location. Furthermore, if this secondary branch happens to split, it mostly splits in two at maximum. Branching or splitting more than twice should remain rare events. Secondly, the graph’s edges are relatively short: **<*w*>** is around 6 µm for a graph diameter ***D*** of 1017 µm. This provides an average estimation of the mean distance between two consecutive branches along tubules or of the mean branch segment length. However, the cumulative edge length ***W*** is over 0.2 m, which is considerable when compared to the total sample volume (3.2×10^-3^ mm^3^) and is indicative of a relatively high linear edge density of tubules and branches (∼ 6.8×10^4^ mm/mm^3^). Third, the largest graph component ***S*** comprises more than half the graph nodes, revealing a strong degree of interconnections. The remaining part of the graph is scattered in about ***K*** ∼ 6000 minor components with size varying from 0– 1% of the total number of nodes. This can at least partly be explained by the finite extent of the image stack, with many isolated nodes or groups of nodes found at or near the boundary which originate from branches or tubules outside the acquisition volume. Finally, the graph seems to have a low resilience as the fault sensitivity ***FS*** is over 0.6, meaning that only a limited number of edges are essential to reach all nodes. The low clustering coefficient (***C*** ∼ 0.02) also confirms a low probability of alternative deviation passing by immediate neighbors of an edge’s endpoints.

Interestingly, while graph cleaning has a massive influence on the number of nodes (−58 %) and edges (−62 %), the impact on other metrics seems rather limited for some and quite critical for others. The main degree remains close to **<*k*>** ∼ 2 because most of the errors were node groups with a degree 1 and a degree 3 (called “spurious edges”, discussed below in III.3.1). Similarly, the total edge length ***W*** remains higher than 2 m and the largest graph component ***S*** still contains > 50 % of graph nodes. This indicates that most of the edges removed were typically spurious edges of short length. As a result, the average length **<*w*>** nearly doubles. Because ***N*** and ***E*** are roughly divided by 2, ***γ*** doubles as it depends linearly on ***E***, but quadratically on ***N***. Also, the increase in ***K*** shows that some erroneous edges were connecting components that shouldn’t be, but this didn’t affect the graph resilience as ***FS*** only slightly decreases.

#### III.2.2 Spatial graph fluctuations over the full volume

To emphasize the spatial changes in network topology as a function of distance to DEJ, we generated coarse-grained maps in Fig.4 and S6 Fig. of the full volume shown in Fig.3.a. All metrics emphasize a topological transition at an approximate distance of 300 µm from the DEJ indicated by a dashed line in Fig.4, although they don’t necessarily exhibit the same contrast between the two areas. For instance, ***N***, ***W***, ***FS*** and **<*w*>** in (Fig 4.b-e) exhibit strong variations. **<*w*>** highlight the small edge length near the DEJ caused by a greater number of branches. Also, shortest path-based metrics such as **<*l*>** (S6 Fig.) appear noisier. This is most likely due to the small size of the sliding window used to calculate the metrics maps (25×25×20.6 µm^3^) which may be insufficient to fully capture path characteristics.

**Figure 4:**
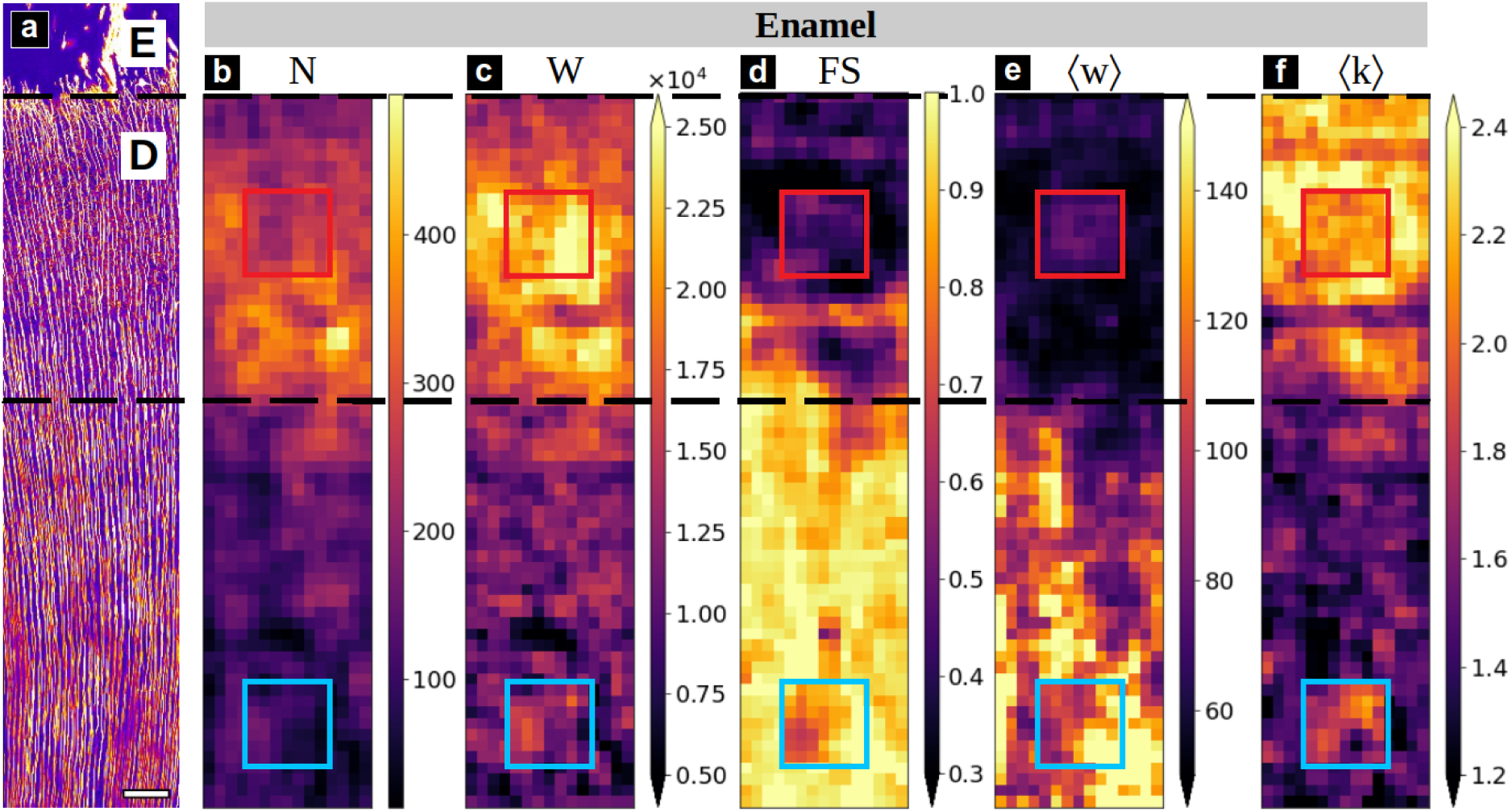
Spatial graph fluctuations. a) Maximum intensity projection of the image stack, shown as a spatial reference. Scale bar 50 µm. b-f) Metric maps of the cleaned graph. b) number of nodes (***N***), c) total edge length (***W***), d) Fault sensitivity (***FS***), e) mean edge length (**<*w*>**) and f) mean degree ( **<*k*>**). Top and bottom squares indicate respectively the ROI 1 and ROI 2, in which values of the manually corrected graph are displayed. Approximate distance from DEJ to transition zone indicated by dashed lines: 300 µm.

Interestingly, a similar transition in mechanical properties was reported by several authors and described in detail by Zaslansky *et al.* as a soft transition zone between enamel and bulk circumpulpal dentin [29]. This gradient was mostly attributed to differences in ultrastructure and mineralization, but our results suggest that the porosity network topology may also contribute to this mechanical gradient. Note that the two topologically different ROIs identified from visual inspection of the confocal data in Fig.3.b and Fig.3.d are also represented in Fig.4 and were found to be representative of the two areas about the 300 µm transition.

#### III.2.3 Ground truth comparison in representative ROIs

The previous observations assume that most graph errors were corrected during the cleaning step. This is typically a non-trivial task that relies on one’s ability to identify artifacts and define sufficiently precise correction rules. To verify the cleaning efficiency, the two topologically distinct regions of interest (ROI) previously identified (Fig.3-5) were manually corrected to create ground truth (GT) graphs.

In addition to the full graph, Table 2 provides a comparative summary of the relative errors between the generated or cleaned graphs and GT for the two ROIs. Overall, the cleaning efficiency is relatively high with respect to the number of nodes and edges, albeit with substantial differences between ROIs. As expected, the initially generated ***N*** and ***E*** are reduced by a factor of ∼ 2 (ROI 1) – 3 (ROI 2) resulting in a 10 % error for ROI 2, which seems reasonable at first sight, while ROI 1 retains a 24 % and 30 % excess ***N*** and ***E***, respectively, which may appear insufficiently precise. However, the relative impact of cleaning is much less pronounced in ROI 2, where all paths and resilience metrics are rarely affected and retain relatively high error rates, while most metrics improve in ROI 1. The resulting cleaned graphs metrics are all within 15 % error of the GT for ROI 1 except the average edge length **<*w*>** (−25 %) and ***FS*** (−33 %), while for ROI 2, the only metrics under 15 % error are **<*w*>**, ***W*** and ***FS***, all others being above 70 % except **<*k*>** (−19 %) and ***γ*** (−31 %) which are somewhat in between. So, overall, the impact of cleaning on connectivity analysis is clearly much more efficient in ROI 1 than ROI 2 where many results may seem unreliable.

**Table 2:** Graph analysis of the two manually corrected ROIs. The values for different metrics are compared between the generated, cleaned and ground truth (GT) graphs.Values between parentheses are the relative differences between the generated or cleaned graph and the GT.

| Metric | ROI 1 |  |  |  |  | ROI 2 |  |  |  |  |
| --- | --- | --- | --- | --- | --- | --- | --- | --- | --- | --- |
|  | Generated |  | Cleaned |  | GT | Generated |  | Cleaned |  | GT |
| $N$ | 8294 | (+185%) | 3683 | (+26%) | 2915 | 3613 | (+259%) | 1187 | (+18%) | 1006 |
| $E$ | 9361 | (+219%) | 4142 | (+42%) | 2925 | 3314 | (+301%) | 790 | (-5%) | 827 |
| $\langle k \rangle$ | 2.29 | (+9%) | 2.33 | (+10%) | 2.11 | 1.86 | (+6%) | 1.42 | (-19%) | 1.75 |
| $\gamma (\cdot 10^4)$ | 2.77 | (-62%) | 6.34 | (-13%) | 7.25 | 5.17 | (-71%) | 11.99 | (-31%) | 17.45 |
| $\langle w \rangle$ | 2.56 | (-63%) | 5.22 | (-25%) | 6.95 | 2.81 | (-74%) | 11.24 | (+5%) | 10.75 |
| $W (\cdot 10^{-3})$ | 24.8 | (+10%) | 23.2 | (+3%) | 22.5 | 9.6 | (-4%) | 10.1 | (+0%) | 10.1 |
| $\langle l \rangle$ | 108 | (-23%) | 122 | (-14%) | 141 | 49 | (-72%) | 44 | (-75%) | 173 |
| $D$ | 289 | (-22%) | 318 | (-14%) | 372 | 144 | (-76%) | 174 | (-72%) | 611 |
| $DI$ | 2.00 | (-20%) | 2.27 | (-9%) | 2.50 | 1.82 | (-45%) | 1.69 | (-49%) | 3.33 |
| $S$ | 0.93 | (+11%) | 0.85 | (+2%) | 0.83 | 0.05 | (-87%) | 0.03 | (-94%) | 0.43 |
| $K$ | 173 | (-23%) | 202 | (-11%) | 226 | 400 | (+94%) | 405 | (+97%) | 206 |
| $FS$ | 0.41 | (-24%) | 0.36 | (-33%) | 0.54 | 0.96 | (+10%) | 0.95 | (+9%) | 0.87 |

The analysis confirms our initial visual perception of a greater vessel density near the DEJ (ROI 1), with higher values for ***N***, ***E*** and ***W***, and higher connectivity shown by higher **<*k*>** (less end points of degree 1, more vessel connections of degree 3) and ***S***, resulting in a lower ***FS***. The high ***FS*** value in ROI 2 emphasizes the overall fragility of the network. The relative size of the largest component ***S*** and the number of connected components ***K*** indicate that the generated and cleaned graphs are heavily fragmented in comparison to the GT in ROI 2 while being very close in ROI 1. This implies that in ROI 2, metrics focusing on this largest component only describe a small part of the topology as apparent in Fig.5. The similar GT values for the number of components ***K*** in both ROIs suggest that the effect of artificial vessels pruning at the imaged volume borders is the same in both cases.

**Figure 5:**
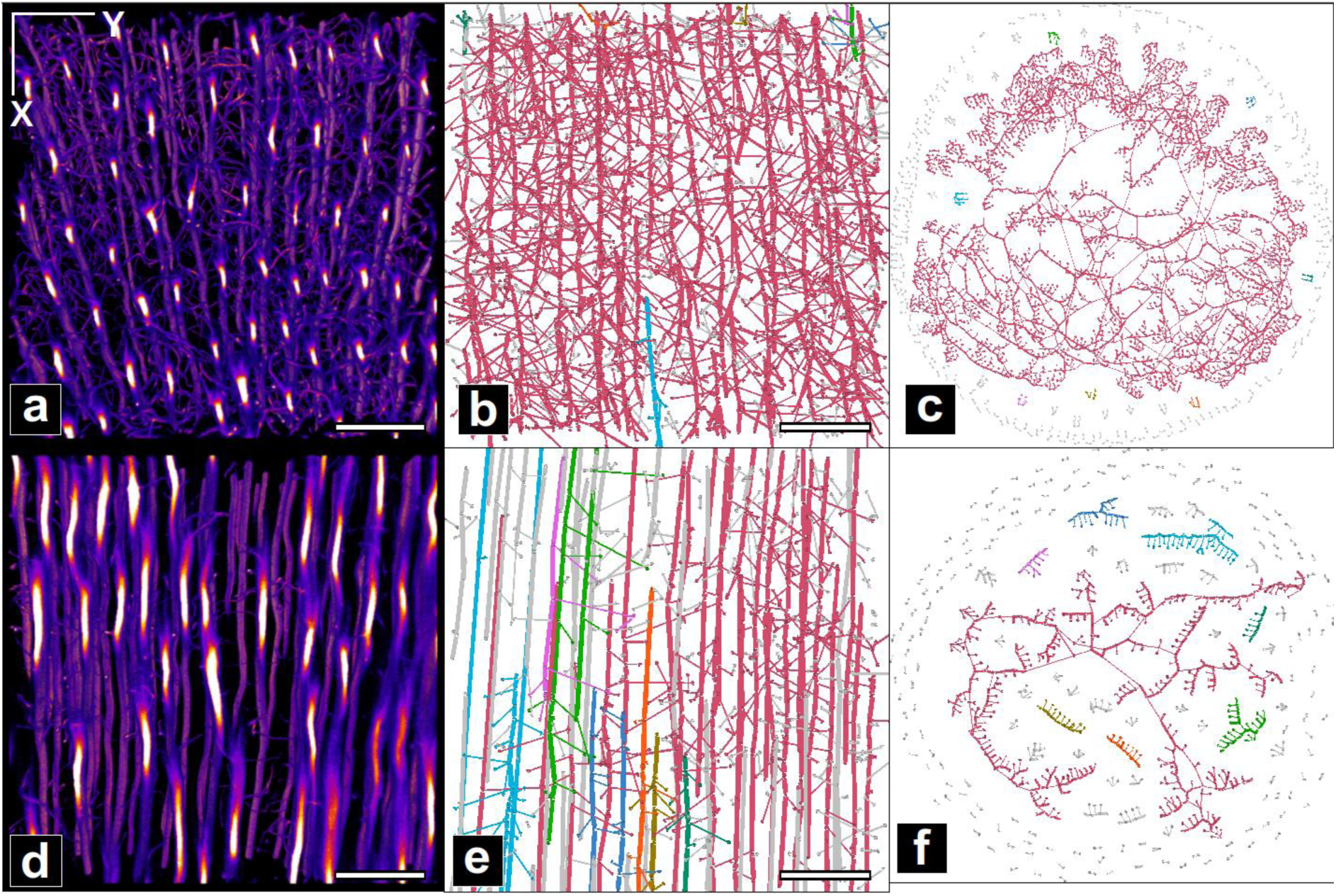
Ground truth graph visualization. a) MIP of ROI 1 region in the original image; scale bar 20 µm. Ground truth graph in ROI 1, shown in b) with spatially localized nodes or in c) with a “Force Atlas” representation (cf II.3.4) highlighting the different components. d), e) f) same representations for ROI 2. Colors indicate the different components, with red associated with the largest one.

Those observations allow better quantifying the important differences in structures in each region. In ROI 2, tubules are connected by a few branches only, thus defining an irregular, sparse 3D grid (Fig.5.f). I ROI 1, branches tend to connect more often, leading to a more complex structure providing multiple alternative paths, which has a direct impact on distance-based metrics (**<*l*>**, ***D*** and ***DI***). For instance, the large ***D*** value in ROI 2 (611.3 µm), which is about 6 times the length of the ROI in X or Y, could be explained by the need to travel further along a given tubule to reach the few available bridges allowing to hop onto another tubule. In ROI 1, ***D*** is smaller because a greater number of bridges are available, thus requiring lesser distance to be traveled along tubules and the same logic goes for **<*l*>** and ***DI***. Nevertheless, it is worthwhile noting that ***DI*** values are rather high in both regions (2.50-3.33), meaning that the transport distance separating two given nodes is 2 to 3 times higher than the Euclidean distance. In comparison, Giacomin *et al.* computed the detour index (or circuity in the study) of metropolitan road networks in the US in [61], and obtained an average value of 1.339 (with a maximum of 1.467. Although these networks are 2D planar, whereas ours are in 3D, the obtained values of 2.5 and 3.33 are indicative of poor transport efficiency.

#### III.2.4 Interpretability of local topological changes

The comparison of the (cleaned) extracted graph for the whole data set to two GTs showed that there are potentially different levels of errors remaining in different ROIs after cleaning (as compared to the GTs). However, the comparison of the two ROIs also reveals quite different topologies in each data set. Our ability to identify topological differences therefore depends on those relative differences. To verify this, we calculated the ratio of the maximum discrepancy between the cleaned graph and the GT (Δ(Cle)) and the difference between the ROIs GT (Δ(GT1/2)) (in Table 3). Under this definition, a ratio < 1 would indicate that the remaining errors in the cleaned graph are sufficiently low to accurately detect topological changes in the whole volume (providing that ROI 1 and 2 represent extreme values), while a ratio > 1 prevents concluding with sufficient certainty.

**Table 3:** Comparison between remaining graph errors and topological differences. *Δ(Gen)* and *Δ(Cle)* are respectively the maximum between the two ROIs of the absolute difference between the generated graph or cleaned graph and the GT. *Δ(GT1/2)* is the absolute difference between ROI 1 and ROI 2. The ratio between *Δ(Gen)* (or *Δ(Cle)*) and *Δ(GT1/2)* indicates how important is the error bias when computing the metric on the generated/cleaned graph in comparison to the ROIs variations (the lower the better).

| Metric | $\Delta(Gen)$ | $\Delta(Cle)$ | $\Delta(GT_{1/2})$ | $\frac{\Delta(Gen)}{\Delta(GT_{1/2})}$ | $\frac{\Delta(Cle)}{\Delta(GT_{1/2})}$ |
| --- | --- | --- | --- | --- | --- |
| $N$ | 5379 | 768 | 1909 | 2.82 | **0.40** |
| $E$ | 6436 | 1217 | 2098 | 3.07 | **0.58** |
| $\langle k \rangle$ | 0.18 | 0.33 | 0.36 | **0.50** | **0.92** |
| $\gamma$ | 12.28 | 5.46 | 10.20 | 1.20 | **0.54** |
| $\langle w \rangle$ | 79.4 | 17.4 | 37.9 | 2.09 | **0.46** |
| $W$ | 2.35 | 0.73 | 12.41 | **0.19** | **0.06** |
| $\langle l \rangle$ | 1247 | 1295 | 321 | 3.89 | 4.04 |
| $D$ | 4673 | 4372 | 2397 | 1.95 | 1.82 |
| $DI$ | 1.51 | 1.64 | 0.83 | 1.82 | 1.98 |
| $S$ | 0.377 | 0.406 | 0.402 | **0.94** | 1.01 |
| $K$ | 194 | 199 | 20 | 9.70 | 9.95 |
| $FS$ | 0.13 | 0.18 | 0.33 | **0.39** | **0.55** |

Table 3 shows that the smallest uncertainty ratio (highest confidence) for the cleaned graph is obtained for the total edge length sum ***W*** (0.06). This ratio is remarkably low compared to those of all other metrics (> 0.37), therefore ***W*** should be considered as a key measure for local analysis. ***N***, ***E***, ***γ***, **<*w*>** and ***FS*** should also prove reliable since their uncertainty ratio is < 0.5–0.6. The mean degree <***k***> will also be considered as its ratio remains < 1 and so could ***S*** which is close to 1, but all other metrics (**<*l*>**, ***D***, ***DI***, ***K***) should be discarded.

This distance from the GT to the cleaned graph can also be visualized in the spatial maps of Fig.4 where the ROI insets represent their GT metric values. While ***N*** and ***W*** are not very different from the surrounding map, the two ROIs clearly appear distinctively for <***k***> in the form of a shift in contrast. This also implies that the fluctuations observed in the images on both sides of the transition zones are most likely significant except for <***k***>.

### III.3. Modeling the sources of errors and assessing their impact

Previous results provide important insights on the metrics sensitivity to remaining errors, allowing to identify which ones are most reliable. However, our analysis is only valid for our imaging setup and graph extraction pipeline and results may vary for other experimental conditions. To assess the generalizability of our observations and better quantify the impact of different parametrization, we computed the evolution of graph metrics under an increasing amount of each type of errors taken independently.

#### III.3.1 Nature and classification of the sources of errors

During the manual correction of the cleaned graphs, repeating error patterns were observed which led to their categorization in four classes: *spurious edges*, *nodes disconnections, missing edges* and *edge bridges*. The characteristic features and type of applied corrections are described in table 4 and Fig.6-8. *Spurious edges* were found to be the most common type of error in the graph, equally distributed between the two ROIs but were almost entirely automatically corrected during the cleaning procedure: 1656 in ROI 1, 1094 in ROI 2, corresponding to a decrease of almost 50% of ***N*** and ***E***. *Nodes disconnections*, on the other hand, couldn’t be entirely corrected during the cleaning step. This was the least common error among the four types but was equally distributed in both ROIs (∼50 in each). *Missing edges* were also not fully corrected in the cleaning step. This was the second most common error during manual correction in ROI 2 (∼100 added edges) but was very rare in ROI 1. On the opposite, *Edge bridges* was the second-most common type of error in ROI 1 but were very scarce in ROI 2: 2190 bridges were detected and removed during the cleaning step (448 in ROI 1, 15 in ROI 2), but an extra 50 % had to be manually corrected in the GT.

**Figure 6:**
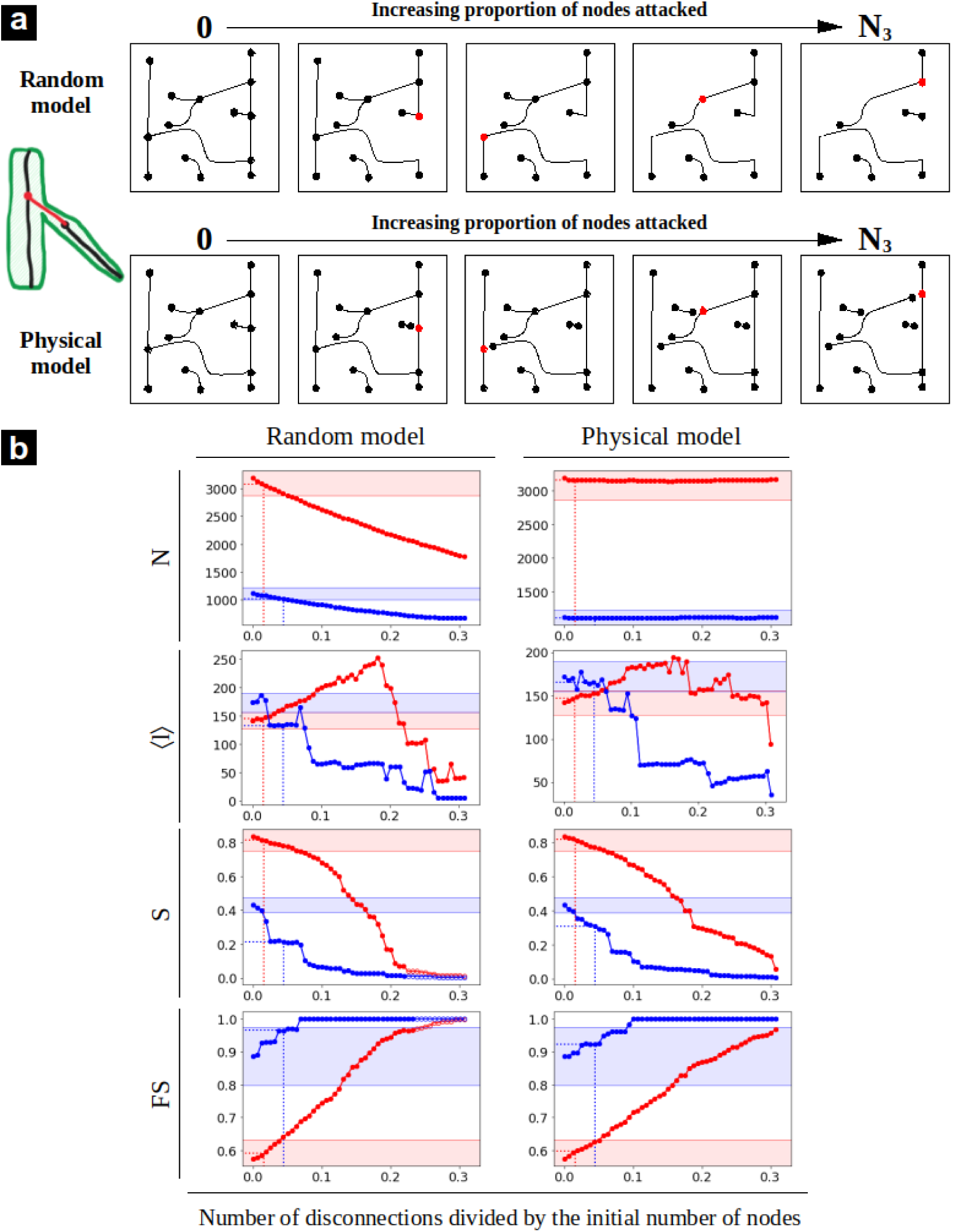
Node disconnections error simulations. a) Schematic representation of the random and physical simulations. b) Simulation results for selected metrics (all results available in S3 Fig.). The red and blue curves correspond respectively to the ROI 1 and 2.

**Table 4:** Characteristics and correction rules of the four classes of identified defects.

| Name | Characteristics | Possible origin | Correction |
| --- | --- | --- | --- |
| **Spurious edges** | The simplest type of artifact, in the form of short edges ($l < 2\mu\text{m}$) with a node of degree 1 stemming from a tubule. | Commonly known artifacts resulting from the skeletonization of rough (pixelated) vessel surfaces [62]. | Removal of the edge and its two nodes (which also fuses two edges on the tubule). |
| **Missing edges** | Correspond to (at least parts of) branches with weak intensity or low contrast in the raw images. | Typical results from insufficient excitation laser power or low fluorescent dye concentration (e.g. small vessel or blocked diffusion). The segmentation parameters may also be another source. | Adding an edge between the existing end nodes. |
| **Node disconnections** | Disconnections which often appears at branch/tubule junctions. | Well known artifacts of vesselness filters which tend to under-perform at intersections [63]. | Adding an edge from the node tip of the disconnecting branch to the tubule edge, followed by the concatenation of the branch and the created edge. |
| **Edge bridges** | Artificial junctions between two crossing non-connecting branches. | Mainly arise from small branches connecting due to the enlargement of tubules or branches by the instrument PSF [16]. Possible enlargements of voxels by filtering are also possible, although less likely. | Removal of the connecting edge followed by the application of the 2-degree nodes correction (as described in II.2.3.) to its ends. |

**Table 5:** relative importance of error type as a function of network topology.

|  | Highly connected region<br>(ROI1) | Less connected region<br>(ROI2) |
| --- | --- | --- |
| Node disconnections | — | — |
| Branch bridges | ++ | — — |
| Missing edges | — — | ++ |

#### III.3.2 Simulation of errors

Using the ground truth in the two ROIs, we simulated two types of error models: *random modifications*, akin to widely performed “*graph attacks*” [56] and targeted “*physics- or biology-informed*” modifications which make use of the spatial graph attributes, as described in II.2. The main difference between the two models essentially resides in their interpretability: random attacks are mostly focused on graph topology by targeting nodes or edges, while the physics-based simulation may alter nodes or parts of edges by thresholding their attributes depending on a physical parameter related to imaging. All results can be found in S3–S5 Appendices Fig.S3-S5 and are summarized in Fig.6-8 below. In order to facilitate the comparison between the random and physical models, additional graphs were added in the middle column of Fig.7.b,8.b representing the correspondence between the physical parameters and the fraction of affected edges.

**Figure 7:**
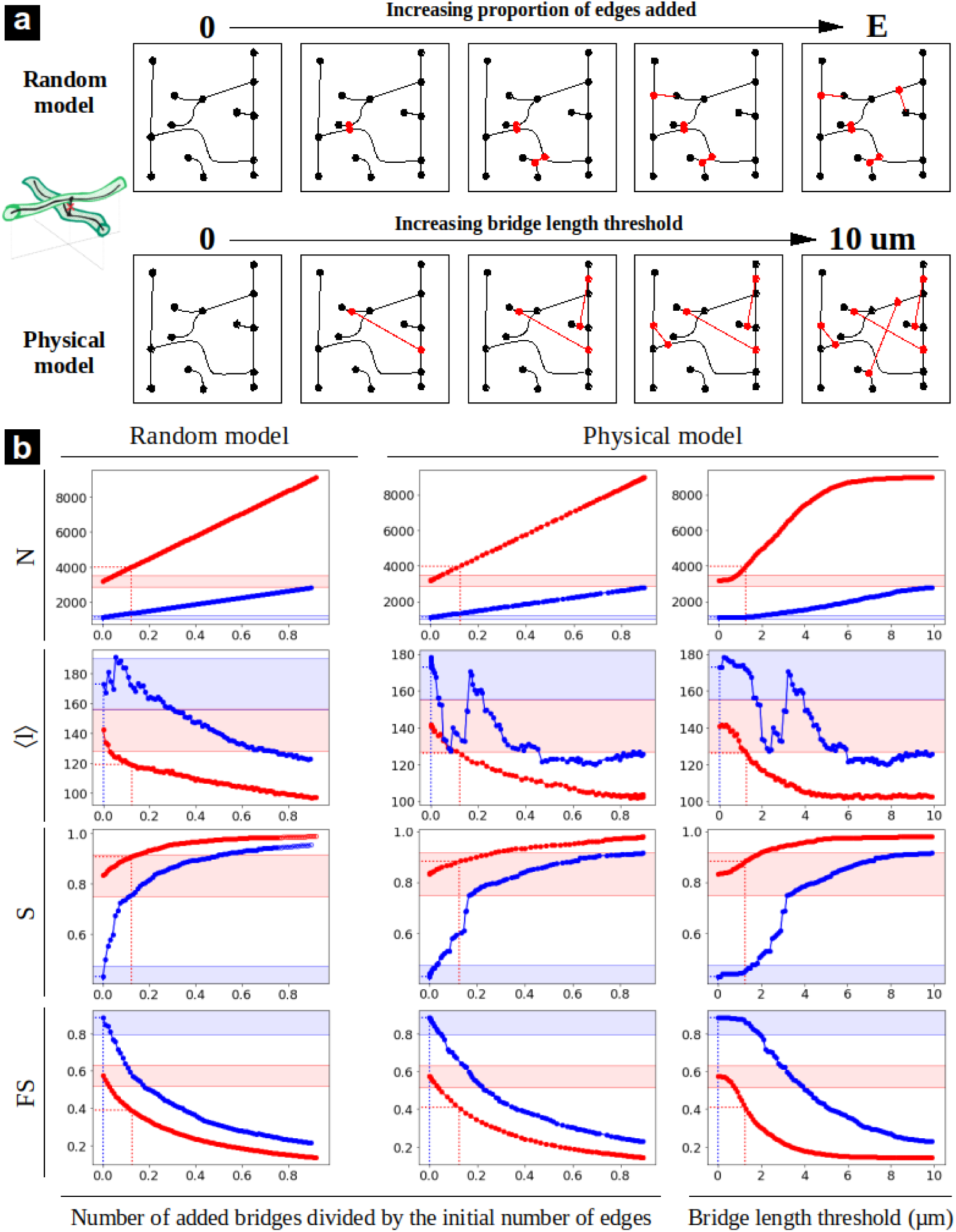
Edge bridges error simulations. a) Schematic representation of the random and physical simulations. b) Simulation results for selected metrics (all results available in S4 Fig.). The red and blue curves correspond respectively to the ROI 1 and 2.

Since *spurious edges* were almost entirely corrected during the cleaning step, we will only consider the three remaining types in the following discussion. From a very general perspective, all results obtained for *nodes disconnections* and *edge bridges* with the random model and those based on physics using the intermediate representation appear to be very close or, at least, follow the same tendencies. A notable exception is <***w***> which follows an inverse trend in both models. In the case of *missing edges*, the two models don’t necessarily yield the same results, so both will have to be examined separately in this case. Also, for all considered type of error and metrics, ROI 1 and 2 curves are distinct but their relative evolution can basically follow three different scenario: 1) they can diverge or follow similar trends but never intersect (e.g. Fig.7.b); 2) they can start with opposite trends and intersect at a point of confusion (black dashed line) which is often found at a relatively low level of error (e.g. **<*l*>** in Fig.6.b,8.b) or 3) they converge after a critical error level beyond which both curves are indistinguishable (e.g. ***S***, ***FS*** in Fig.6.b,8.b). However, each type of error simulation exhibits different patterns which must be described separately:

##### Node disconnections

this simulation (Fig.6 and S3 Fig.) assesses the impact of improper branching connections, a typical artifact of the vesselness filter. The ***random model*** simply fully deletes a fraction of edges randomly, so ***N*** and ***E*** decrease linearly. The ***physical model*** only disconnects a portion of nodes with degree 3, starting with the smallest diameter branches, such that ***N*** and ***E*** remain constant up to the maximum fraction of affected nodes (30 %) corresponding to the total number of nodes of degree 3 found in the graph. At the end, nodes of degree 0 are removed, and edges connected to degree 2 nodes are concatenated and the node removed as well. As a result, <***k***> naturally drops while the average edge length <***w***> increases in both models due to the edges being merged. Naturally, the total edge length ***W*** decreases.

This type of defect evidently results in a quick degradation of the graph integrity with a very high fragmentation (***S*** < 0.1) for only ∼10 % of edges removed in the least connected region (ROI 2) and 20-30 % for the most connected one (ROI 1) as shown in Fig.5.b. Interestingly, this leads to very different behaviors of path metrics in the two regions: in the least connected one (ROI 2), the average shortest paths **<*l*>** rapidly decreases as parts of the main component get disconnected (decreasing ***S***). In the most connected region (ROI 1), **<*l*>** increases due to longer detours needed to overcome the disconnections, as long as the main component still contains ***S*** ∼ 40 % of edges. It reaches a peak after the removal of ∼ 20% of the graph nodes after which it drops. ***D*** logically follows a similar trend and so does ***DI***, although the reason for this is less clear.

Unlike the physical model, the random model doesn’t target thinner vessels and is more likely to remove tubules. For that reason, the connectivity metrics ***S*** and ***FS*** are affected at much lower values in ROI 1 (Fig.6.b). This emphasizes the fact that tubules are more important than branches for the connectivity in ROI 1.

##### Edge bridges

this simulation (Fig.7 and S4 Fig.) models connections between branches that were often corrected manually in the ground truth. Connections are either added between edges chosen randomly (***random model***) or if their relative minimum distance is smaller than a threshold parameter (***physical model***). The fundamental difference between the two models is that the first can connect edges at long distances while the second connects edges at short, increasing distances. This results in opposite trends for the mean edge length **<*w*>**, but all other metrics exhibit the same behavior in the two models.

Because of the high spatial edge density in ROI 1, ***N*** and ***E*** increase more rapidly than in ROI 2. Consequently, most metrics rise (<***k***>, ***W***, ***S***) or drop (***γ***, **<*l*>**, ***D***, ***DI***, ***K***, ***FS***) more steeply in ROI 1 than ROI 2. Interestingly, all remain virtually unaffected below ∼ 1 µm, indicating that typical distances between vessels are at least greater than this value. Also, all values seem to remain stable beyond ∼ 7 µm in ROI 1 and ∼ 9-10 µm in ROI 2, indicating a typical maximum distance between tubules.

##### Missing edges

this simulation (Fig.8 and S5 Fig.) aims to emulate branches which are visible in the data but appear fragmented in the segmented data, thus also fragmenting the graph. It yields the strongest differences between the random and physical models. The ***random model*** is close to this used for edge disconnections (regardless of their degree in this case), so the results follow the same trends within the 0-30 % equivalent range. Removing fractions of edges based on their minimum image intensity attribute as done in the ***physical model***, on the other hand, yields a different behavior. ***N*** and ***E*** exhibit no change until a critical intensity of ∼ 25, then increase sharply to a maximum of 50 for ROI 2 and 75 for ROI 1 after which they decrease exponentially. This can be understood by the fact that the weakest signal from the branches mostly lie within the 25-50 intensity range, after which, most branches (60 % of edges) have been removed and tubules progressively disappear. Because of the highest edge density in ROI 1, the effect is more pronounced in this region.

**Figure 8:**
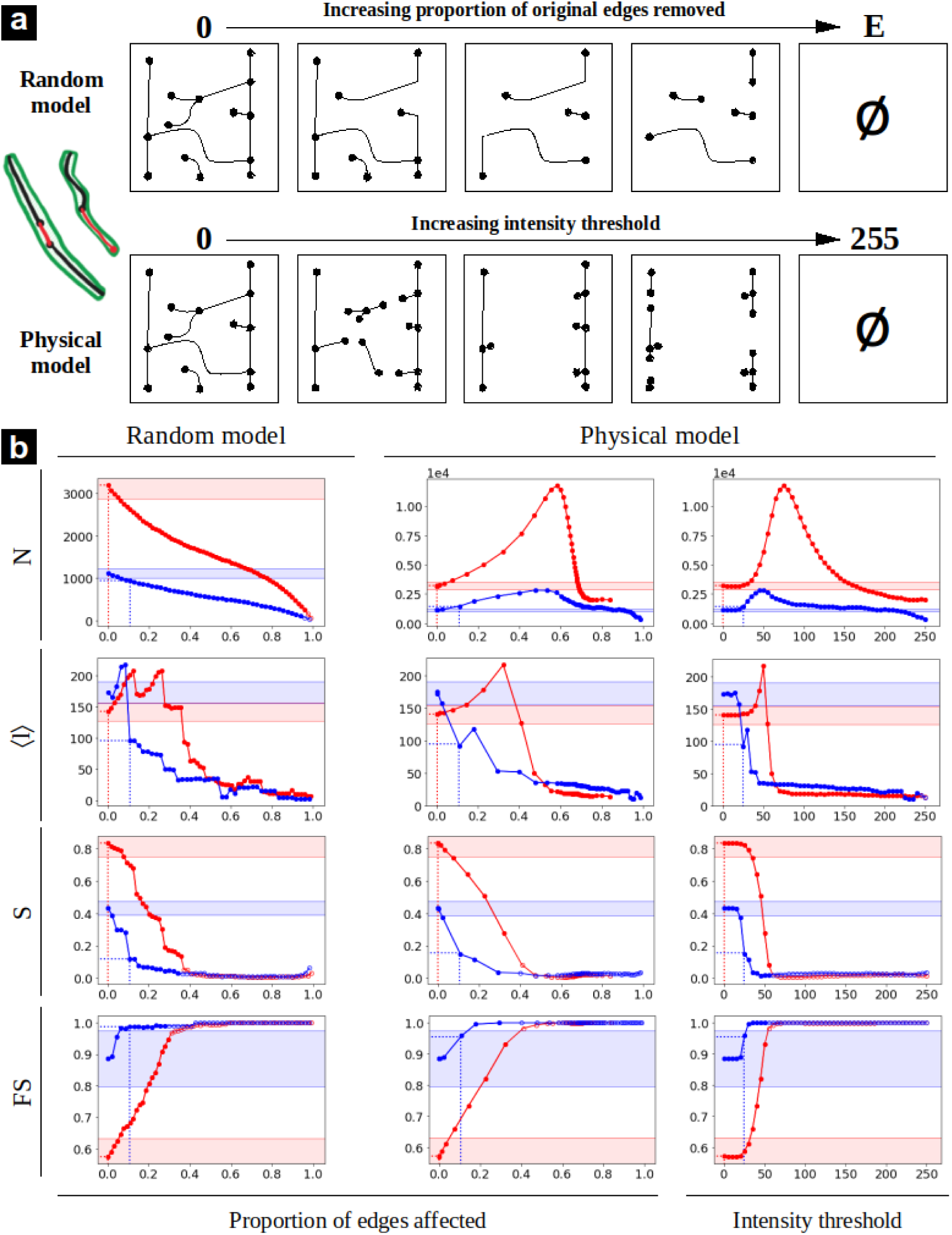
Missing edges error simulations. a) Schematic representation of the random and physical simulations. b) Simulation results for selected metrics (all results available in S5 Fig.). The red and blue curves correspond respectively to the ROI 1 and 2.

In all graph simulations (Fig.6.b-8.b), the first value on the y axis always corresponds to the GT and the dashed lines show the amount of errors reported during the manual correction (x axis), as well as the associated error value on the cleaned graph (Err_CG_) (y axis). In addition, the region corresponding to a 10 % error from the GT value is indicated by shaded areas to highlight an acceptable error rate. This value is somewhat arbitrary, but approximately corresponds to an upper limit of the minimum percentage error between the cleaned graph and the ground truth in ROI 1 and ROI 2 (Table 2) for the metrics which expressiveness was found to be < 1 (Table 3).

All Err_CG_ values for nodes disconnections fall within this 10 % error range except for ***D*** and ***S*** in ROI 2 (Fig.6.b). More importantly, those values are always lower than the confusion points of the metrics (at least for the physical model). Altogether, this indicates that our cleaning graph pipeline for this level of imaging quality is sufficiently robust against nodes disconnection errors. The situation is more delicate for *branch bridges*, where Err_CG_ for **<*l*>** in ROI 1 is located beyond the confusion point in the physical model representation. Furthermore, Err_CG_ values mostly fall on the initial plateau region of ROI 2, but not in ROI 1. Since the metrics are very sensitive to this type of error (sharp increase or decrease following the initial plateau), a slight increase in bridge length (physical model) or fraction of affected edges (random model) results in a significant change in metric value. So clearly, all metrics are highly sensitive to this type of error, which is more problematic in densely connected regions. The situation is reversed for *missing edges*, for which all metrics fall on a stable initial plateau within 10 % error rates for ROI 1 and out of range for ROI 2 (Fig.8.b). More problematic, some Err_CG_ values calculated for ROI 2 (**<*l*>**, ***D***) are beyond the point of confusion, such that their interpretation is opposite to that made from GT values. Therefore, this error is clearly more critical for less connected regions.

## IV. Discussion

Our first aim, in this study, was to show that dentin cellular porosity indeed forms a complex spatial network. In this healthy 23-year old female molar, we observed two topologically distinct regions within the first 300 µm from the DEJ and beyond. A denser branching area was observed close to the DEJ, which corroborates previous SEM and optical imaging studies. At further distances, fewer branches were still observed which proved important for the global network connectivity. Those simple histological differences are reflected by the most basic graph characteristics, such as branching type (node degree **<*k*>**), inter-distance or length (mean edge length **<*w*>**) which all clearly highlight this transition. Such transition at approximately 300 µm cannot be related to mantle dentin which is generally less than 50 µm thick in human molars, but rather coincide with reported variations in collagen fibrils organization and mineralization [29]. We therefore hypothesize that the gradient in mechanical properties (e.g. hardness, Young modulus etc.) could at least partly also be accounted for by the distribution of dentinal porosity, as previously suggested [12].

However, our graph analysis provides a much clearer view of the overall network connectivity: e.g., 83 % of nodes were found to be connected in the main component of ROI 1, compared to 43 % in ROI 2 (***S*** in table 2), as illustrated in Fig.5. Those differences are highly significant and the fact that both regions are strongly interconnected is, as such, an important result which tends to indicate that tubules cannot be considered independently from a functional point of view. This observed porosity may or may not contain odontoblastic processes (not visualized in our study). If they do, the observed interconnections directly represent a path to inter-cellular connectivity evidenced by TEM [6]. Otherwise, they are simply filled with physiological fluid, in which case fluid flow through these interconnections could favor collective odontoblasts activation through shear stresses according to the main mechanosensing theory [1]. Either way, the observed topological connectivity should have a direct consequence on mechanosensing which should be different between the two regions, in accordance with the known increase in dentinal sensitivity in coronal dentin close to the DEJ.

The total edge length ***W*** is also a very important parameter in spatial graphs as it provides a direct measure of the “cost”, in a general sense, associated with the network [55]. For cellular networks, it represents the metabolic cost of maintaining the cells alive and connected. In mineralized tissues, maintaining the pericellular space (or residual tubular porosity if the main process has retracted) also requires additional cellular activity to prevent unwanted tissue formation and mineralization [64]. Again, we observed a marked difference between the two regions, with only half the value in ROI 2 compared to ROI 1 (0.101 and 0.22 m respectively in Table 2). The fault sensitivity ***FS*** also provides a useful quantification of how fragile the network is, should a connection (tubule or branch) disappear. It is therefore inversely proportional to the resilience of the network. With values close to 0.4 close to the DEJ and 0.9 beyond 300 µm, the network strength appears very different in the two zones (Fig.7): quite resilient close to the DEJ and more fragile after the transition zone. Because the network resilience is a critical factor for long-lived cells, such analysis clearly emphasizes the usefulness of graph analysis.

Although the analysis of a single sample prevents drawing definite biological conclusions, this pilot study allowed examining a basic set of common classes of graph metrics and testing their sensitivity in the specific context of dentin cellular porosity networks. In doing so, we thoroughly analyzed the impact of potential artifacts that may arise from confocal fluorescence microscopy or subsequent image processing to extract the network graph. A first important result is that certain types of metrics were found to be highly prone to residual errors in the graph. This is the case for nearly all path metrics, **<*l*>**, ***D*** and ***DI***, which are particularly sensitive to edge bridges and missing edges, with typical values of final cleaned graph beyond the point of confusion, which basically results in opposite interpretation of the relative values in ROI 1 and 2 compared to the ground truth (Fig.7-8). This is somewhat problematic since those metrics are important quantities to understand the efficiency of transport in graphs.

Our simulations of the relative importance of different types of errors based on physical parameters allows a much finer interpretation of the impact of errors in possibly very different experimental conditions than reported for this study. For instance, *branch bridges* essentially characterize the impossibility to distinguish neighboring branches or tubules that may not be physically connected but appear so due to a lack of imaging resolution or improper segmenting with low intensity thresholding. The optical point-spread-function (PSF) used in our study was typically ∼ 300 nm in the imaging plane and ∼ 700 nm axially. This yielded a typical value of branching in our graphs of 1.2 µm which would correspond to the distance between two branches of 500 nm diameter separated by the axial PSF. This is observed to have a strong effect on all metrics of ROI 1, which is denser and thus more prone to such artifacts. But this PSF is actually obtained in close to optimal conditions, with a close index matching between sample and mounting media and high-quality oil objective of 1.3 NA. So, improving these measuring conditions would be difficult with a standard confocal microscopy setup and possibly require super-resolution microscopy, which may prove very tedious. In fact, since many measurements of cellular networks in mineralized tissues are often performed in PBS with water objectives of much lower NA, we anticipate that many results could be obtained with worse performance than ours. A simple increase in axial PSF of 700 nm to 1 µm would result in a ∼ 10 % apparent increase of connectivity ***S*** in ROI 1 and 2 or an equivalent loss in fault sensitivity.

The case of *missing edges* is potentially even worse. Such artifacts typically result from imperfect tubule or branch fluorescent staining due to lack of dye diffusion in remote or partly occluded cellular porosity, yielding very low intensity values in the acquired images. Imperfect segmentation is unavoidable in such cases. Such loss of information is very often observed in published studies of the lacuno-canalicular network in bone (see e.g., [34,43]). In regions of low connection density, this may easily result in a 25-50 % error or higher on virtually all graph metrics (e.g. with a threshold of 30-50 on ROI 2 in Fig.8). It should be noted here that this reasoning is not limited to confocal fluorescence microscopy. Resolution limitations and low signal/background ratio is an inherent problem to any imaging modality, although each would have its own set of advantages/disadvantages. For example, X-ray CT has isotropic resolution, which is a strong advantage over optical methods and may decrease edge bridges with equivalent in-plane resolution. But this would require achieving spatial resolutions in the ∼ 100 nm range which has not yet been demonstrated with laboratory sources for dentin imaging. Our own experience shows that a proper imaging of dentinal branches with X-ray CT requires advanced synchrotron phase contrast imaging, which cannot be put on the same level as widespread confocal microscopy.

Similarly, image processing remains another major area for improvement. For instance, should the PSF be accurately estimated (which is difficult with strongly aberrating mineralized tissues), image deconvolution could probably reduce the amount of *edge bridges*. Another non-negligible source of error are *nodes disconnections*. Those are a very well-known artifact of vesselness filtering which tends to under-perform in branching areas [63]. Although we used the best known version of this filter, more advanced correction rules could be applied, e.g. as geodesic voting which was successfully implemented by Peyrin *et al.* for bone studies [65]. Supervised or unsupervised deep-learning models could also prove useful for such purposes, but because they would require user annotation or input, those methods would still require benchmarking.

To our knowledge, such detailed error estimation was rarely published before, at least for cellular networks in mineralized tissues. Our study not only shows that this is absolutely mandatory to discuss any observed topological fluctuation which could otherwise be at the statistical noise level, but also that it provides quite deep insights into experimental and image processing aspects that need to be considered and might be improved. The result of our error modeling could thus be used by a wide community of microscopists to estimate the amount of errors or the degree of precision of graph analysis of dental porosity performed in different imaging conditions, providing the characteristics of the instruments be known a priori.

## V. Conclusion

In this study, we investigated the topology of cellular porosity in crown dentin of a young female adult molar and showed that it exhibits characteristic features of a complex network even at millimetric distances from the dentin-enamel junction. We defined nodes as connections between tubules surrounding the main odontoblastic process and lateral branches formed by thinner secondary processes and edges as portions of those processes. We were thus able to show that a very large fraction of odontoblasts in the measured volume could be interconnected to some degree. This could have strong biological consequences in terms of cell-cell interactions, spatial extent of stimulated zones under localized mechanical/temperature stresses that may induce a more complex mechanosensing response than currently envisaged in clinical practice. Based on the examination of basic network features, such as node degree or edge lengths, we were also able to quantify the transition between the well-known highly branched region in the immediate vicinity of the DEJ from this further away. Interestingly, all more advanced network characteristics such as shortest paths or node fraction in the main connected component were found to be sensitive to this topological transition area. Moreover, the network resilience was found to strongly differ in the two regions. However, our in-depth evaluation of potential artifacts arising from the experimental image acquisition and post-processing showed that some graph metrics were very sensitive to such bias. This particularly concerns metrics measuring possible paths in the network, which ultimately relate to possible information transfer pathways between cells. We further classified the nature of encountered errors that couldn’t be fully corrected automatically during image processing and simulated their specific impact on the network characteristics. Our modeling also provides a useful framework for other experimental conditions that may be encountered in confocal microscopy studies and image processing of cellular networks in dentin or other mineralized tissues and, possibly, other types of networks.

## Supporting information

**S1 Fig: Graph cleaning algorithm.** a) Graph cleaning pipeline. b) Detailed description of the branch bridges removal conditions (all must be fulfilled).

**S2 Fig: Degree distribution of the different graphs used in the study.** Hatched parts correspond to nodes of degree 1 that are close to the border, indicative of vessels being cut by the borders of the acquisition.

**S3 Fig: Node disconnections error simulations:** A) Simulation results for metrics N, E, 〈k〉, γ, 〈w〉 and W. B) Simulation results for metrics 〈l〉, D, DI, S, K and FS. The red and blue curves correspond respectively to the ROI 1 and 2.

**S4 Fig: Edge bridges error simulations:** A) Simulation results for metrics N, E, 〈k〉, γ, 〈w〉 and W. B) Simulation results for metrics 〈l〉, D, DI, S, K and FS. The red and blue curves correspond respectively to the ROI 1 and 2.

**S5 Fig: Missing edges error simulations:** A) Simulation results for metrics N, E, 〈k〉, γ, 〈w〉 and W. B) Simulation results for metrics 〈l〉, D, DI, S, K and FS. The red and blue curves correspond respectively to the ROI 1 and 2.

**S6 Fig: Spatial maps of network metrics.** a) number of edges (E), b) edge density (γ), c) mean shortest path (<l>), d) diameter (D), e) detour index (DI), f) relative size of the largest component (S), g) number of connected components (K)

## Acknowledgements

We thank Prof. Ariane Bernal (Sorbonne Univ, Univ Paris Cite, Ctr Rech Cordeliers, Inserm, U1138, Equipe Physiopathol Orale Mol, Paris) for valuable discussions concerning odontoblast development.

## Author Contributions

L. Chatelain: Investigation, Data Curation, Formal Analysis, Software, Visualization, Writing – original draft.

N. Tremblay: Supervision, Software, Validation, Writing – review & editing.

E. Vennat: Resources, Validation, Writing – review & editing.

E. Dursun: Resources, Validation, Writing – review & editing.

D. Rousseau: Conceptualization, Methodology, Supervision, Writing – review & editing.

A. Gourrier: Project administration, Funding acquisition, Conceptualization, Methodology, Resources, Supervision, Software, Writing – review & editing.

## Notes

https://doi.org/10.5281/zenodo.14726443

### Competing Interest Statement

The authors have declared no competing interest.

### Summary of Updates

Modified text for clarity: introduction, figure order and tables. Precisions added concerning the potential impact of the work.

### Funding

- International Human Frontier Science Program Organization: RGP0023/2021

