## Supplementary material for "Cellular porosity in dentin exhibits complex network characteristics with spatio-temporal fluctuations": Supplemetary data

#### S1 Appendix: Graph cleaning algorithm

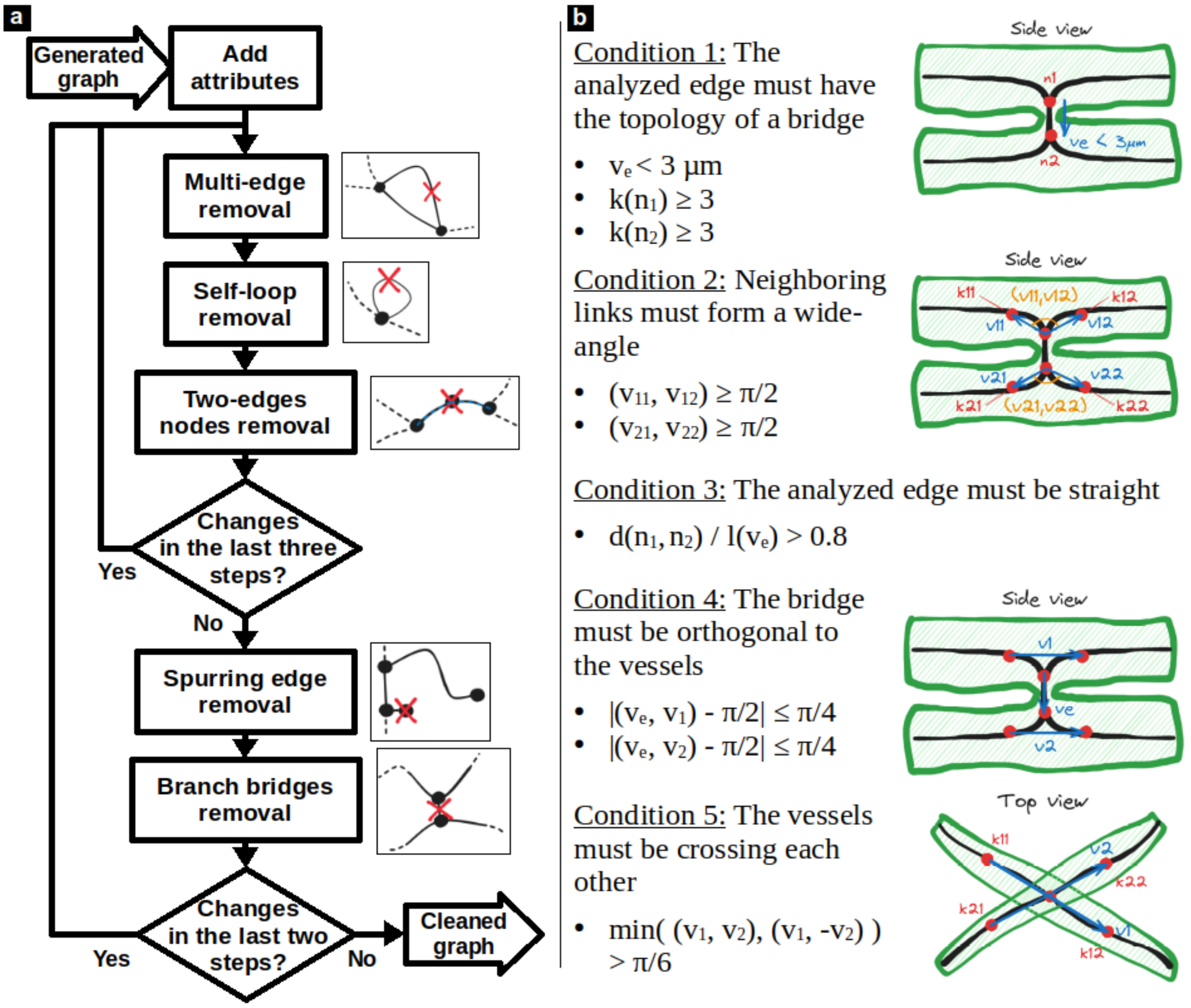


**Fig.S1: Schematics explanation of the graph cleaning procedure.** a) Graph cleaning pipeline. b) Detailed description of the branch bridges removal conditions (all must be fulfilled).

### S2 Appendix: Degree distribution of generated, cleaned and ground-truth graphs

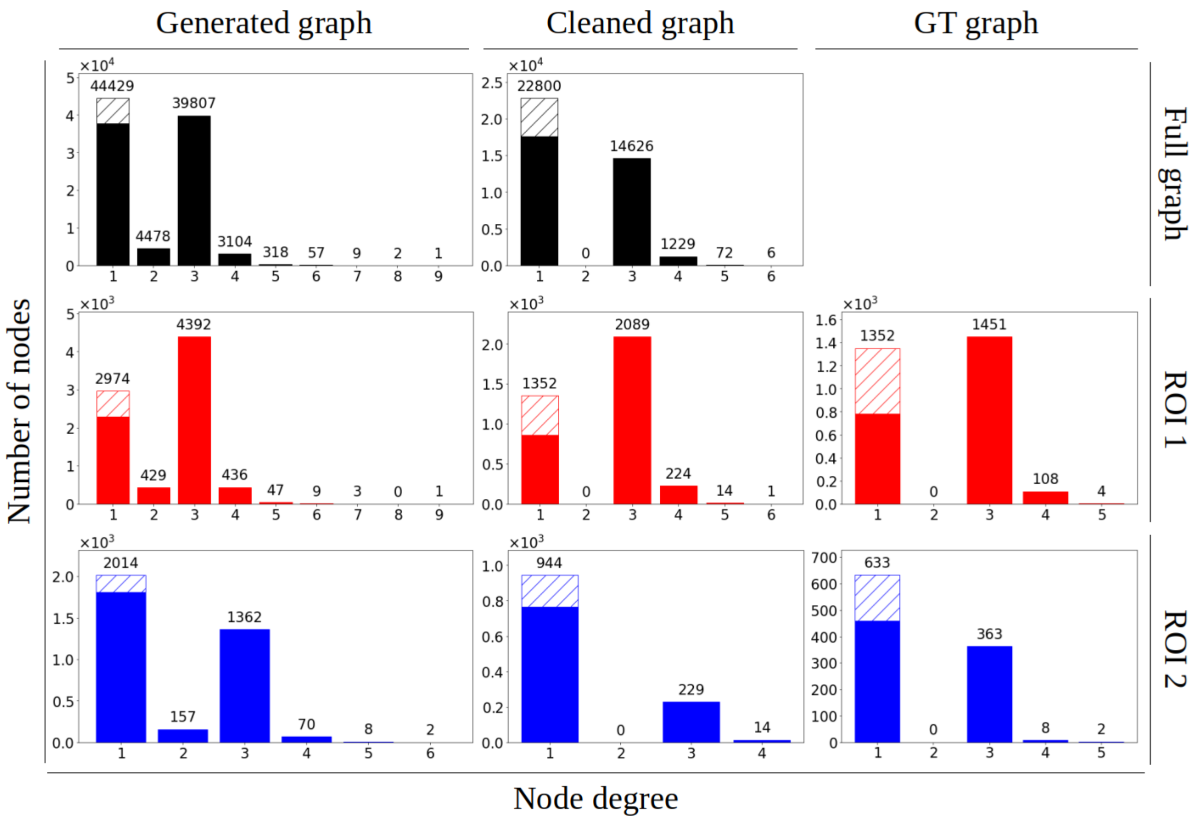


**Fig.S2: Degree distribution of the different graphs used in the study.** Hatched parts correspond to nodes of degree 1 that are close to the border, indicative of vessels being cut by the borders of the acquisition.

### S3 Appendix: Full results for the node disconnections error simulations.

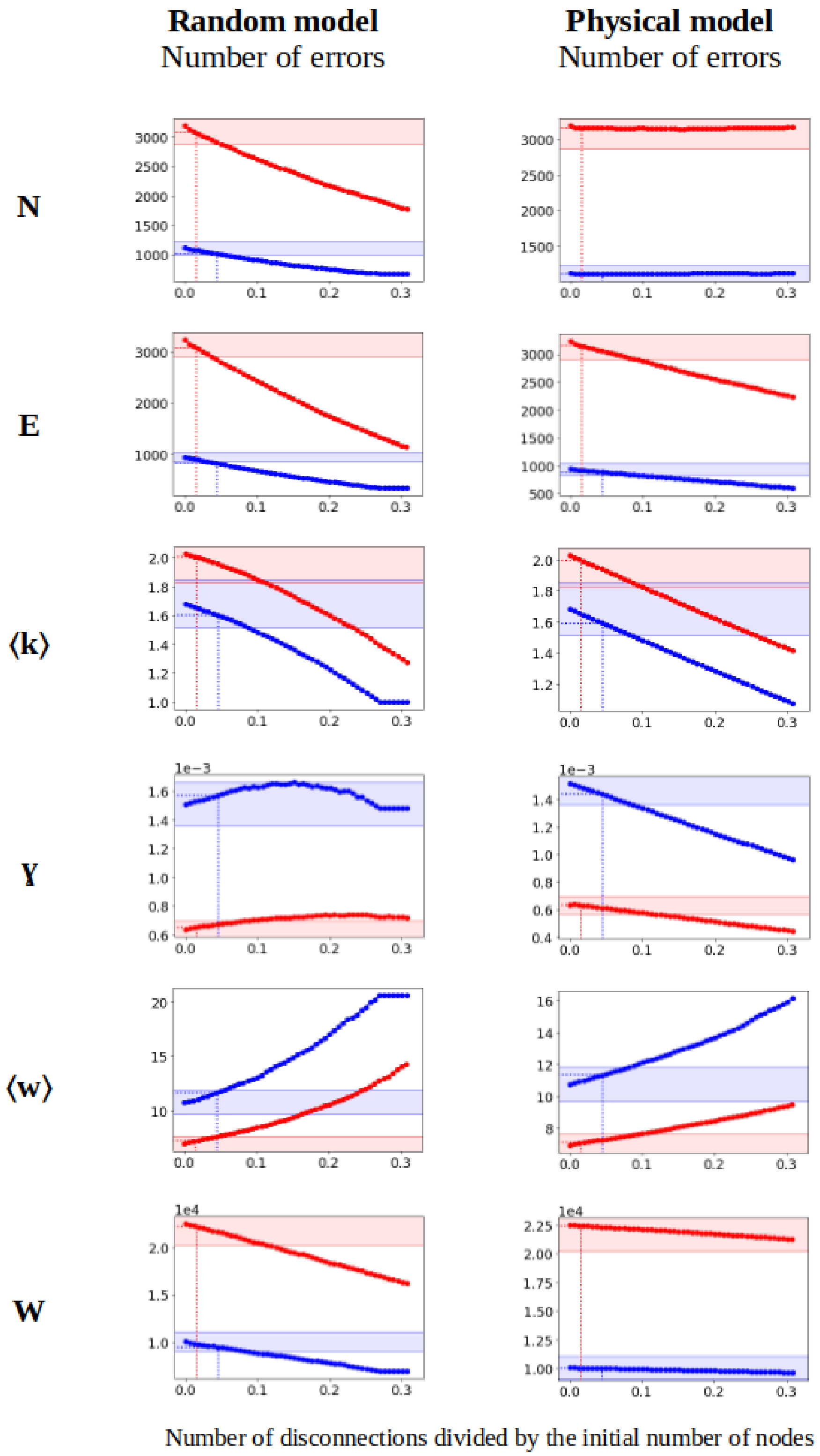


**Fig.S3a: Simulation results for metrics *N* (number of nodes), *E* (number of edges,) ⟨*k*⟩ (mean degree), *γ* (gamma index), ⟨*w*⟩ (mean edge length) and *W* (total edge length).** The red and blue curves correspond respectively to the ROI 1 and 2.

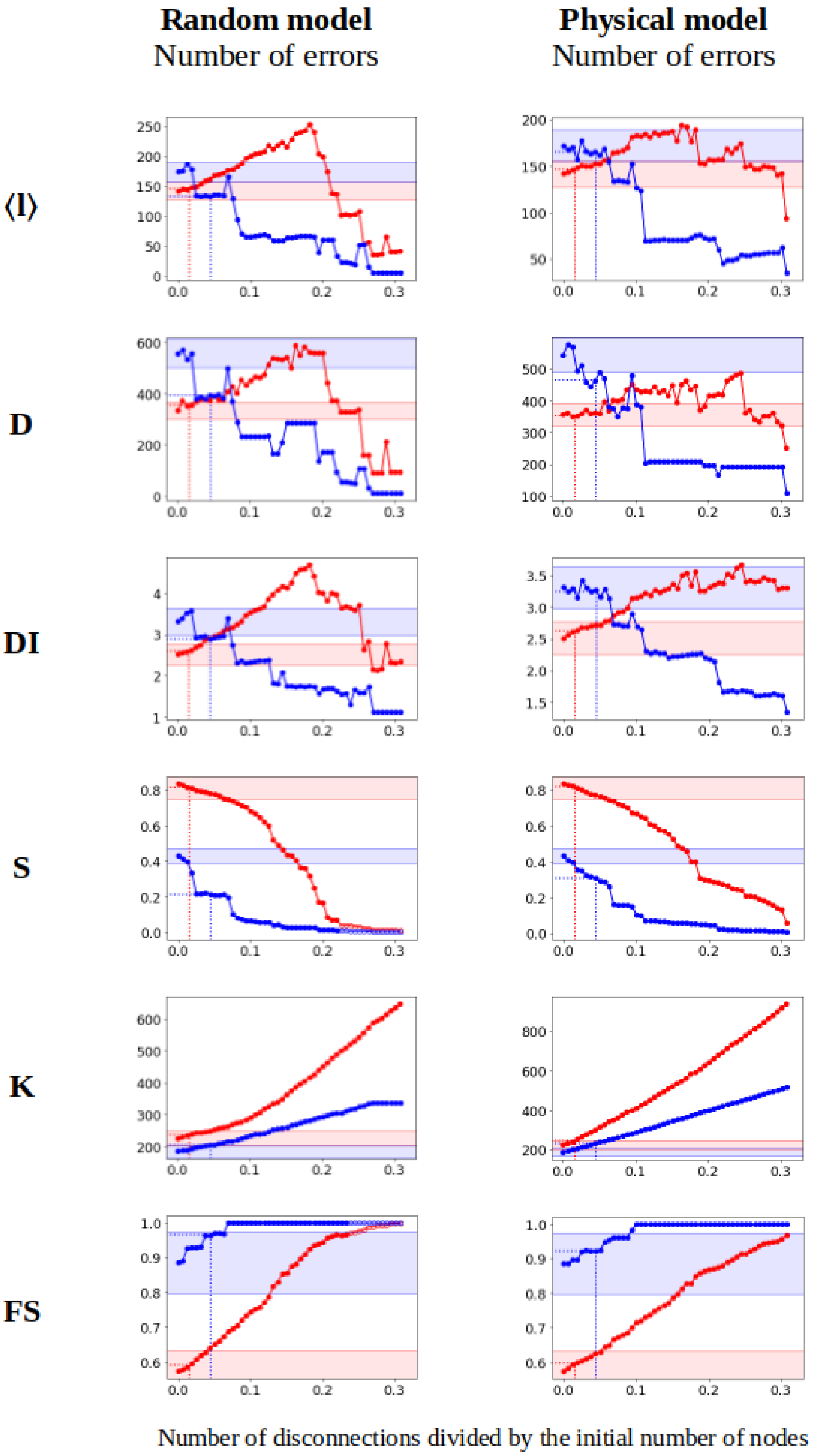


**Fig.S3b: Simulation results for metrics ⟨ l⟩ (mean shortest path), *D* (diameter) *DI* (detour index), *S* (relative size of the largest component), *K* (number of connected components) and *FS* (fault sensitivity).** The red and blue curves correspond respectively to the ROI 1 and 2.

### S4 Appendix: Full results for the edge bridges error simulations.

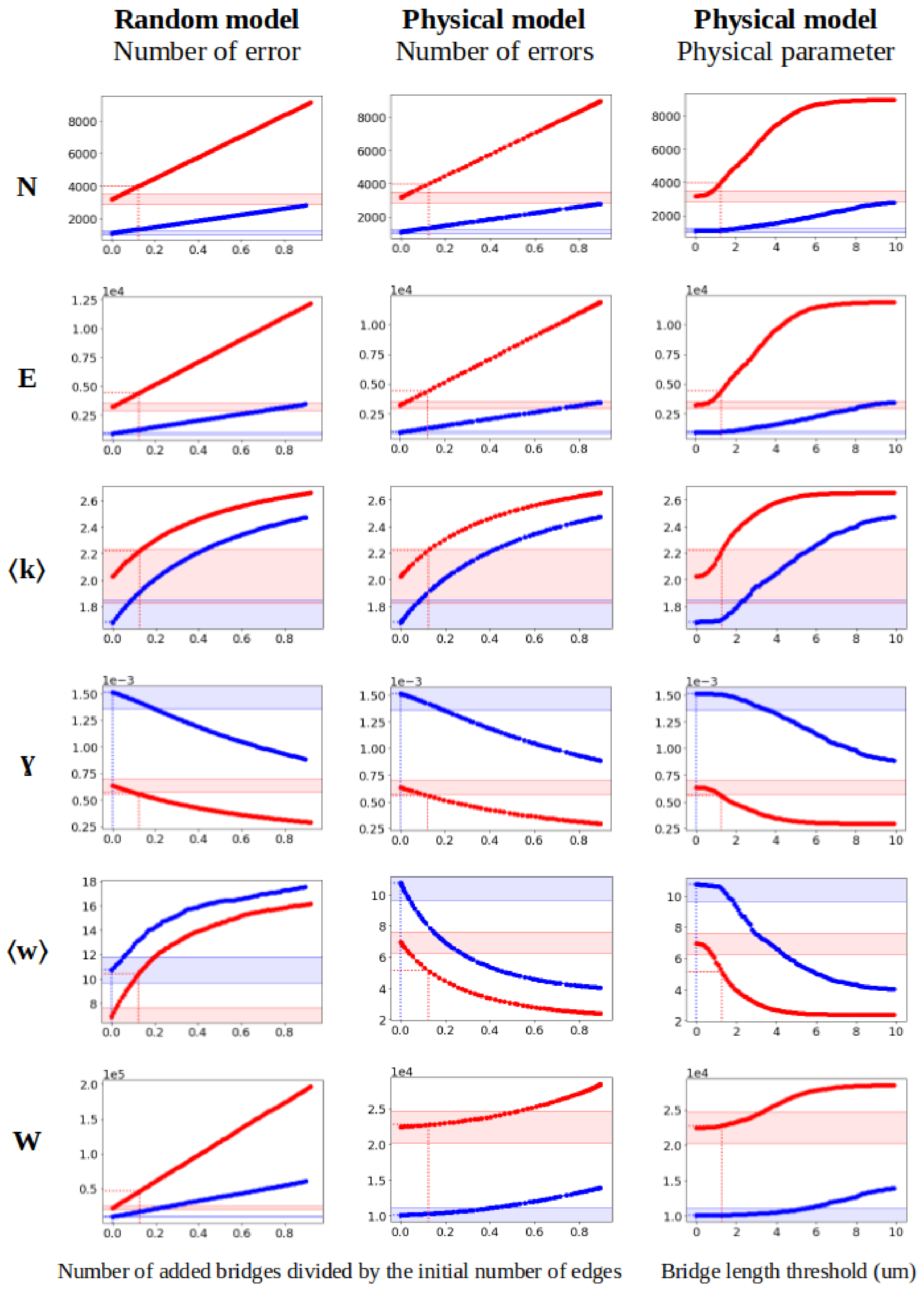


**Fig.S4a: Simulation results for metrics *N* (number of nodes), *E* (number of edges,) ⟨*k*⟩ (mean degree), *γ* (gamma index), ⟨*w*⟩ (mean edge length) and *W* (total edge length).** The red and blue curves correspond respectively to the ROI 1 and 2.

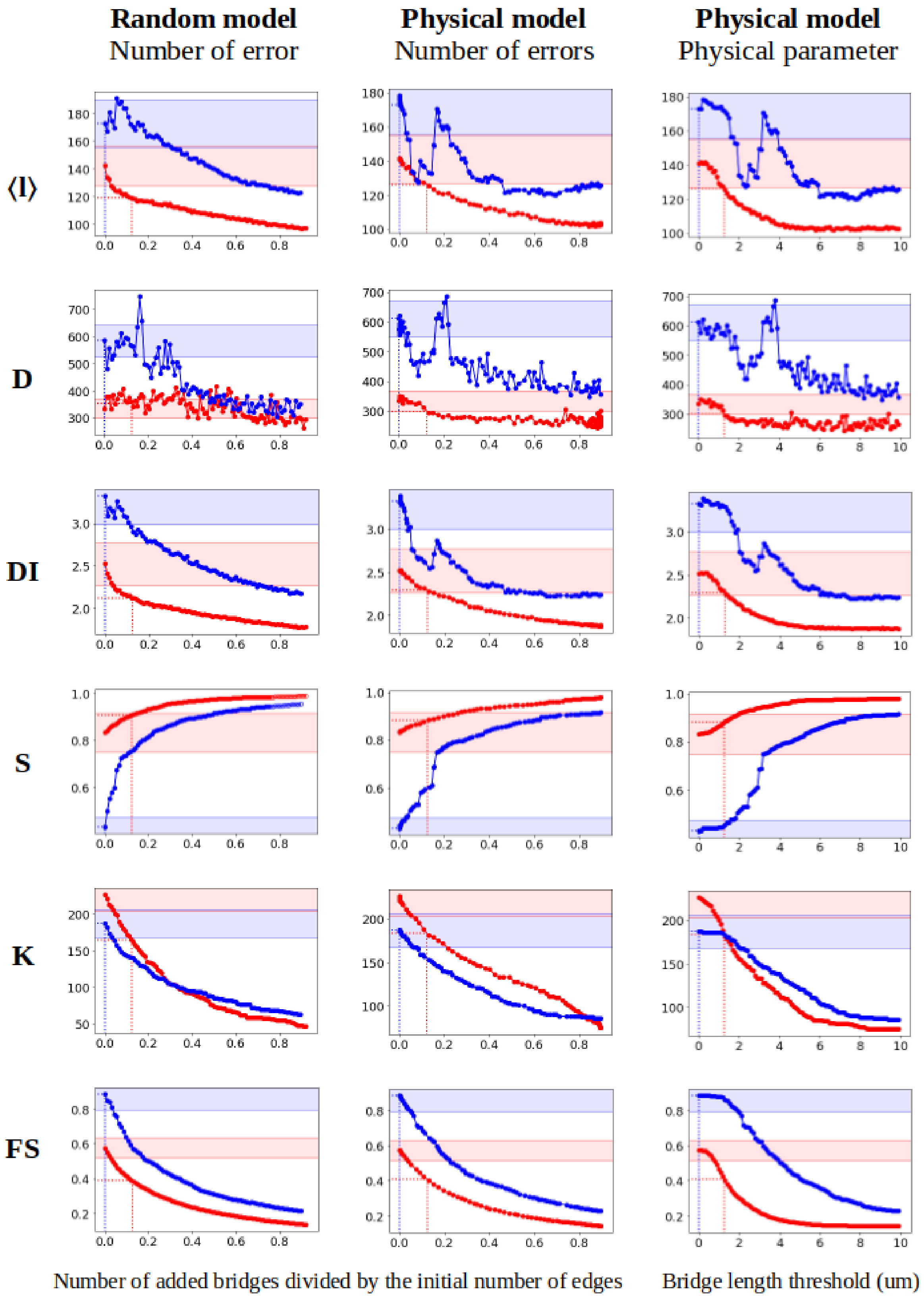


**Fig.S4b: Simulation results for metrics *⟨l⟩* (mean shortest path), *D* (diameter) *DI* (detour index), *S* (relative size of the largest component), *K* (number of connected components) and *FS* (fault sensitivity).** The red and blue curves correspond respectively to the ROI 1 and 2.

### S5 Appendix: Full results for the missing edges error simulations.

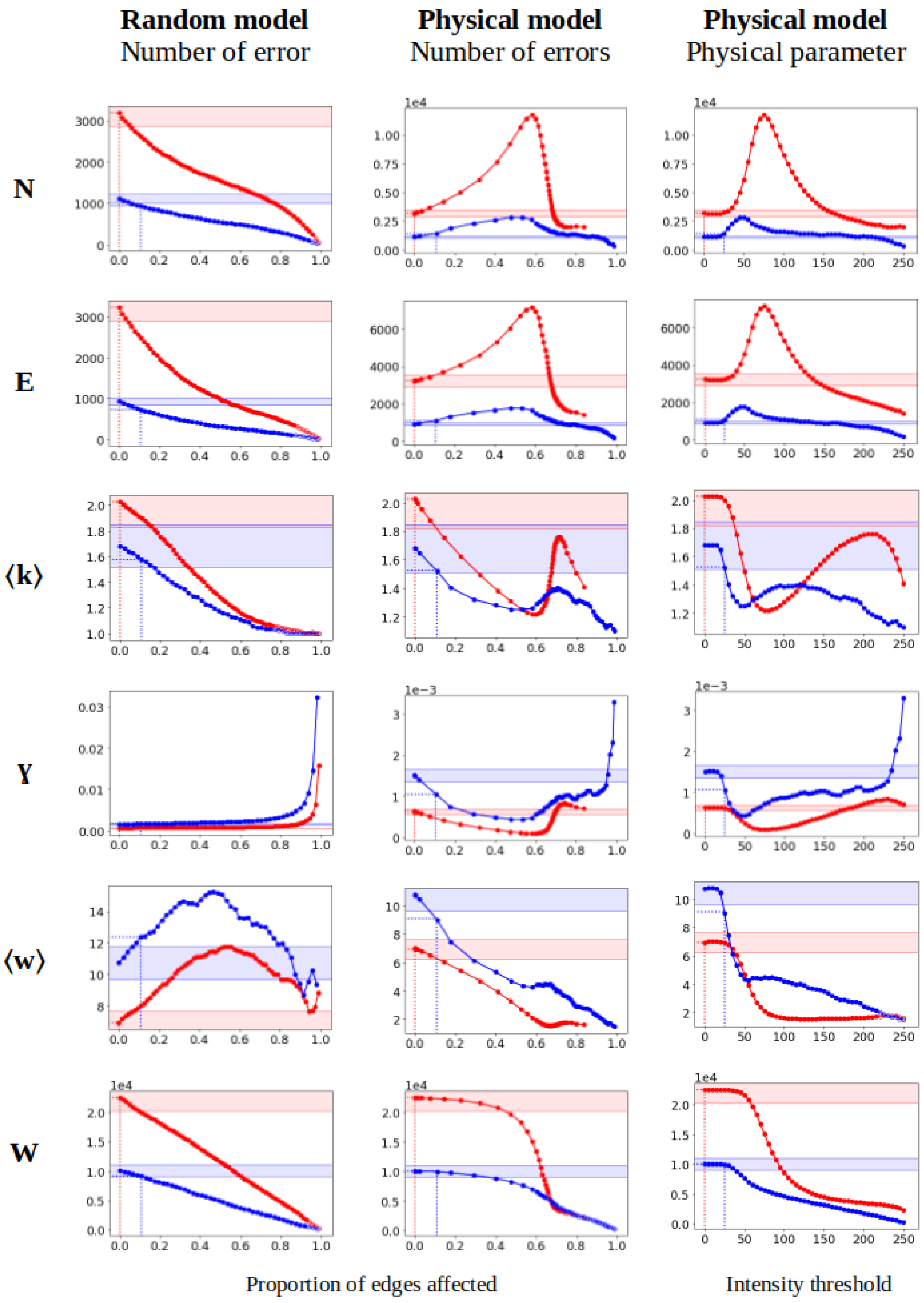


**Fig.S5a: Simulation results for metrics *N* (number of nodes), *E* (number of edges,) ⟨*k*⟩ (mean degree), *γ* (gamma index), ⟨*w*⟩ (mean edge length) and *W* (total edge length).** The red and blue curves correspond respectively to the ROI 1 and 2.

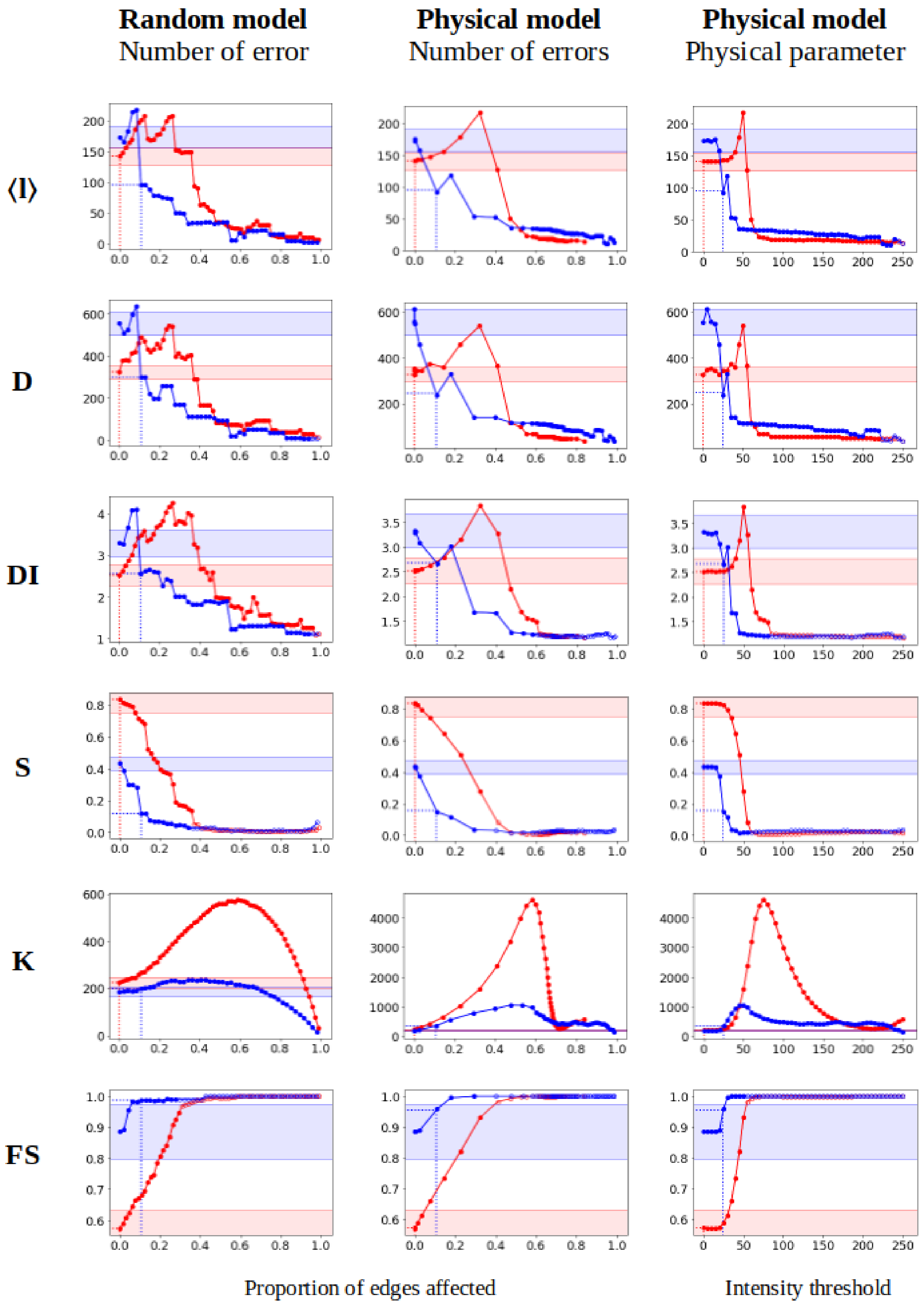


**Fig.S5b: Simulation results for metrics *⟨l⟩* (mean shortest path), *D* (diameter) *DI* (detour index), *S* (relative size of the largest component), *K* (number of connected components) and *FS* (fault sensitivity).** The red and blue curves correspond respectively to the ROI 1 and 2.

### S6 Appendix: Metric maps results

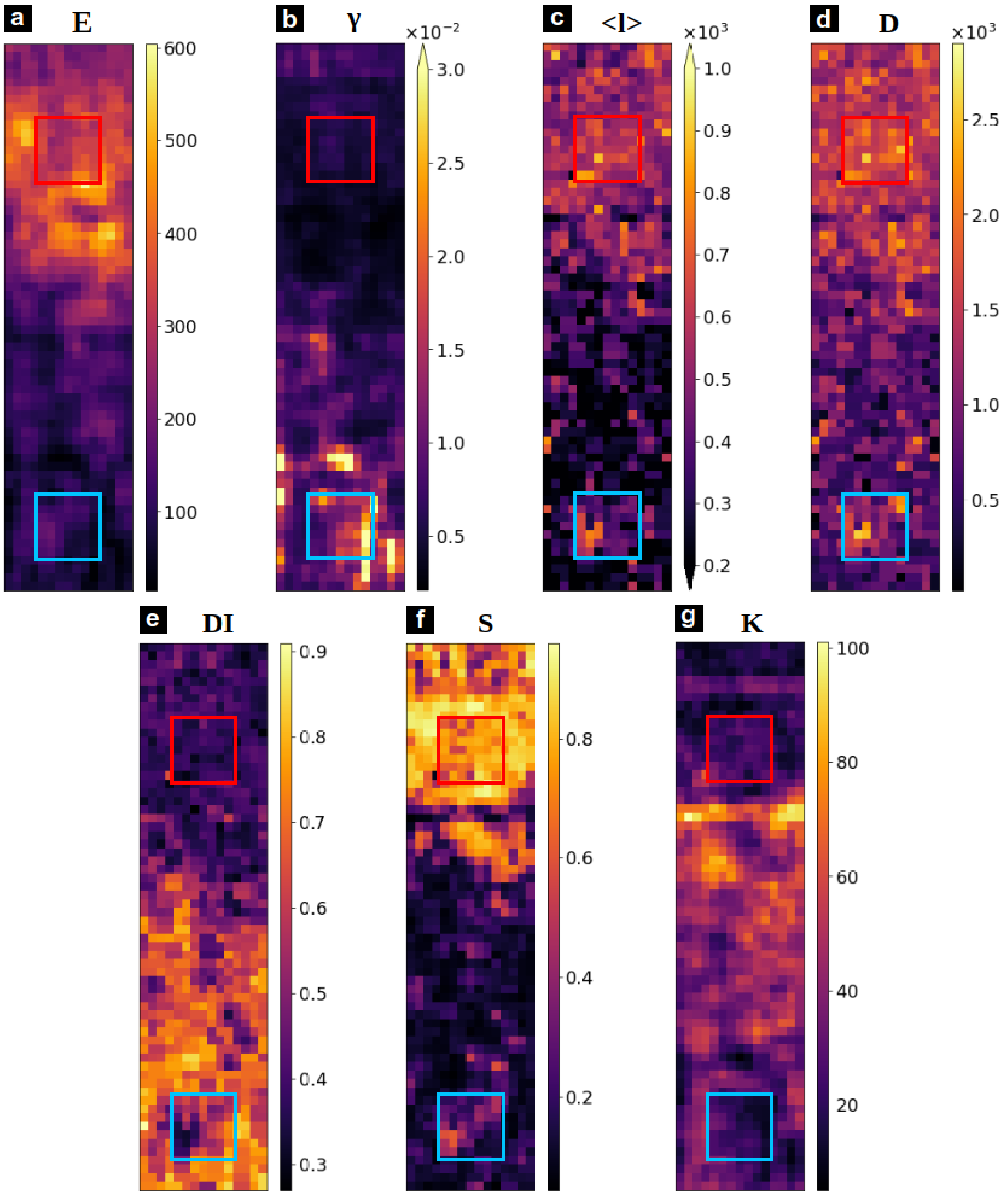


**Fig.S6: Spatial maps of network metrics.** a) number of edges (***E***), b) edge density (***γ***), c) mean shortest path (***<l>***), d) diameter (***D***), e) detour index (***DI***), f) relative size of the largest component (***S***), g) number of connected components (***K***)
